# Dual-compartment engagement of STAR-family proteins SAM68 and QKI by LINC00941 sustains oncogenic fitness in RAS-driven lung cancer

**DOI:** 10.64898/2026.05.11.722569

**Authors:** Disha Acharya, Jean C-Y Tien, Anu Sharma, Vinita Bhat, Aaditya Singh, Mohammad Azharuddin, Sethuramasundaram Pitchiaya, Xuhong Cao, Brendan A Veeneman, Saravana M Dhanasekaran, Bal Krishna Chaube, Arul M Chinnaiyan, Sudhanshu Shukla

## Abstract

Long non-coding RNAs (lncRNAs) are increasingly recognised as effectors of oncogenic signalling, yet the transcriptional programmes through which driver mutations regulate lncRNA expression remain poorly defined. Here we identify LINC00941 as a direct transcriptional target of FOSL1, an AP-1 transcription factor downstream of the KRAS-MAPK pathway, establishing the first FOSL1-regulated lncRNA in lung adenocarcinoma (LUAD). LINC00941 is significantly upregulated in LUAD across multiple independent cohorts, and its depletion via siRNAs, shRNAs, and antisense oligonucleotides (ASOs) induces proliferative arrest and stress-induced premature senescence, accompanied by transcriptomic suppression of cell cycle and DNA damage response (DDR) genes. Mechanistically, LINC00941 operates through a dual-compartment mechanism engaging two STAR-family RNA-binding proteins in distinct subcellular contexts. In the nucleus, LINC00941 binds SAM68 through its 700–1300 nucleotide region and shields it from proteasome-mediated degradation, thereby sustaining SAM68-dependent PARP1 activation and DDR competency; RNF123 is identified as a candidate E3 ligase mediating SAM68 turnover in the absence of LINC00941. In the cytoplasm, LINC00941 sequesters QKI, preventing its nuclear translocation; LINC00941 depletion releases QKI to the nucleus, driving alternative splicing dysregulation including validated NUMB exon 12 exclusion, and QKI co-depletion rescues the anti-proliferative phenotype both in vitro and in xenograft models. Multi-cohort survival analysis across three independent LUAD datasets (n=649) identifies LINC00941 as an independent prognostic factor for poor overall survival. Gymnotic ASO-mediated targeting of LINC00941 significantly suppresses xenograft tumour growth without systemic toxicity, providing preclinical proof-of-concept for therapeutic tractability. Together, these findings establish LINC00941 as a compartment-specific oncogenic scaffold within the KRAS-FOSL1 transcriptional axis and a tractable therapeutic target in LUAD.

## Introduction

Lung cancer is the leading cause of cancer-related mortality worldwide, accounting for an estimated 1.8 million deaths annually and surpassing the combined mortality of colorectal, breast, and prostate cancers [1]. Non-small cell lung cancer (NSCLC) comprises approximately 85% of all lung cancer cases, with lung adenocarcinoma (LUAD) being the predominant histological subtype. Despite the transformative clinical impact of immune checkpoint blockade and molecularly targeted therapies, the overall five-year survival rate for LUAD remains dismal at approximately 28%, underscoring the continued urgency to define the molecular programmes that drive disease progression and to identify tractable therapeutic targets [2]. Oncogenic mutation of KRAS is the most prevalent genomic driver in NSCLC, occurring in 25–30% of lung adenocarcinomas, and constitutively activates the RAF–MEK–ERK mitogen-activated protein kinase (MAPK) cascade to enforce sustained proliferative signalling and promote therapy resistance [3]. Although allele-selective KRAS^G12C^ inhibitors have entered clinical practice, response rates remain partial and acquired resistance emerges invariably, motivating systematic characterisation of the downstream transcriptional programmes that propagate KRAS oncogenic output [4].

Among the transcriptional effectors activated downstream of the RAS–MAPK cascade, MYC and FOSL1 (encoding FRA-1) represent the two principal oncogenic transcription factors that execute the KRAS-driven gene expression programme [5, 6]. The MYC-regulated long non-coding RNA (lncRNA) landscape has been systematically characterised as a core component of the MYC oncogenic programme — most notably KIMAT1, a KRAS-responsive lncRNA that sustains lung tumour growth by reprogramming miRNA biogenesis [7, 8]. In contrast, the FOSL1 arm of this pathway remains entirely unexplored with respect to lncRNA targets. FOSL1, a stringently MAPK-regulated AP-1 family transcription factor that heterodimerises with JUN-family partners to bind AP-1 consensus elements, is consistently overexpressed in KRAS-driven cancers and governs programmes of proliferation, invasion, and epithelial-to-mesenchymal transition [5]. This represents a fundamental and unaddressed gap in our understanding of the KRAS–MAPK transcriptional network: whereas the MYC arm has been extensively linked to lncRNA regulation, no lncRNA has been demonstrated to be a direct transcriptional target of FOSL1 in any cancer context.

Long non-coding RNAs—transcripts exceeding 200 nucleotides with negligible protein-coding capacity—have emerged as critical functional nodes within cancer-associated regulatory networks [9]. A defining and mechanistically consequential property of lncRNAs is their subcellular localisation: depending on whether they reside in the nucleus, cytoplasm, or both, lncRNAs engage distinct molecular machineries and execute fundamentally different functions [10]. Nuclear lncRNAs predominantly operate through epigenetic and transcriptional mechanisms, recruiting chromatin-remodelling complexes, modulating transcription factor occupancy, and regulating nuclear architecture at specialised sub-compartments such as nuclear speckles (e.g., MALAT1) and paraspeckles (e.g., NEAT1) [11, 12]. In contrast, cytoplasmic lncRNAs regulate post-transcriptional processes including mRNA stability and translation, microRNA sequestration as competing endogenous RNAs (ceRNAs), and RNA-binding protein (RBP) activity [10]. Critically, a subset of lncRNAs exhibits dual nucleo-cytoplasmic distribution and can perform compartment-specific and mechanistically distinct functions within the same cell, with some dynamically redistributing between compartments in response to cellular signals [10]. Understanding subcellular localisation is therefore not merely a descriptive characteristic but a prerequisite for delineating the molecular mechanism and biological function of any given lncRNA.

LINC00941 (long intergenic non-coding RNA 00941)—also referred to as lncIAPF (lncRNA in Aggressive Phenotype)—is a chromosome 12p11.21-encoded lncRNA that is upregulated across multiple solid tumour types, including NSCLC, pancreatic adenocarcinoma, gastric cancer, and colorectal cancer, where it promotes oncogenic phenotypes through diverse mechanisms including ceRNA activity, chromatin looping, and protein stabilisation [13]. In NSCLC specifically, LINC00941 has been reported to sponge miR-877-3p to upregulate VEGFA expression, and has been identified as an autophagy-suppressing lncRNA with independent early diagnostic potential in lung adenocarcinoma [14, 15]. Critically, transcriptome-wide comparison of patients with aggressive LUAD (defined by early disease recurrence) against those with non-aggressive disease (long-term recurrence-free survival) identified LINC00941 as the most significantly upregulated transcript in the aggressive cohort, positioning it as a candidate driver of disease severity rather than simply of malignant transformation. Notwithstanding these associations, four fundamental questions remain unaddressed: (i) the upstream transcriptional signals governing LINC00941 expression in aggressive NSCLC; (ii) its precise subcellular distribution; (iii) the compartment-specific protein interactions through which it operates; and (iv) whether it represents a viable therapeutic target amenable to clinically translatable intervention.

Here we address all four of these gaps and establish LINC00941 as a central pathogenic node, a high-fidelity prognostic biomarker, and a druggable oncogene in aggressive LUAD. We demonstrate that LINC00941 is the first lncRNA identified as a direct transcriptional target of FOSL1 downstream of the KRAS–MAPK signalling axis, with FOSL1 occupying the LINC00941 promoter in a MAPK-dependent manner, as established by chromatin immunoprecipitation, luciferase reporter assays, and pharmacological MEK inhibition. LINC00941 exhibits dual nucleo-cytoplasmic localisation and engages a unique, dichotomous mechanism of action: in the nucleus, it acts as a ‘protein shield,’ binding to and stabilising the RNA-binding protein SAM68 (KHDRBS1) to protect it from proteasomal degradation and extend its functional half-life; concurrently, in the cytoplasm, it functions as a ‘molecular anchor,’ sequestering the tumour-suppressive splicing factor Quaking (QKI) to prevent its nuclear translocation and restrict its function. This dual compartment-specific mechanism cooperatively promotes unchecked proliferation while enabling cancer cells to evade oncogene-induced senescence. Functionally, LINC00941 depletion inhibits tumour cell proliferation, induces G2/M cell cycle arrest, triggers oncogene-associated senescence, and suppresses tumour growth *in vivo*. Importantly, inhibition of LINC00941 with therapeutic-grade antisense oligonucleotides (ASOs) significantly suppresses tumour growth *in vivo*, providing preclinical proof-of-concept for its therapeutic tractability. These findings identify LINC00941 as the FOSL1-regulated lncRNA and reveal it as a compartment-specific, multifunctional effector of the KRAS transcriptional network with significant therapeutic relevance in aggressive NSCLC.

## Results

### LINC00941 is transcriptionally upregulated by RAS–MAPK signalling via FOSL1 in lung adenocarcinoma

The KRAS-MAPK signalling axis is a central oncogenic driver in lung adenocarcinoma (LUAD) (Figure 1A); however, its regulation of long non-coding RNAs remains poorly characterised. To identify lncRNAs regulated by this pathway, we performed transcriptomic analysis of the A549 cells (KRAS-mutant) following pharmacological inhibition of MEK/ERK signalling. Genes downregulated upon MEK inhibitor treatment were overlapped with genes overexpressed in KRAS-mutant relative to KRAS-wild-type cells. This integrative analysis, visualised as a starburst plot, identified LINC00941 among the most significantly KRAS-upregulated lncRNAs that were concurrently suppressed by MEK inhibition (Figure 1B–D). Concordantly, pharmacological MEK inhibition in H358 and H1299 cells resulted in a significant reduction in LINC00941 transcript levels in both cell lines as measured by qRT-PCR (Supplementary Figure 1A), further confirming that LINC00941 expression is maintained by active MEK–ERK signalling.

**Figure 1.**
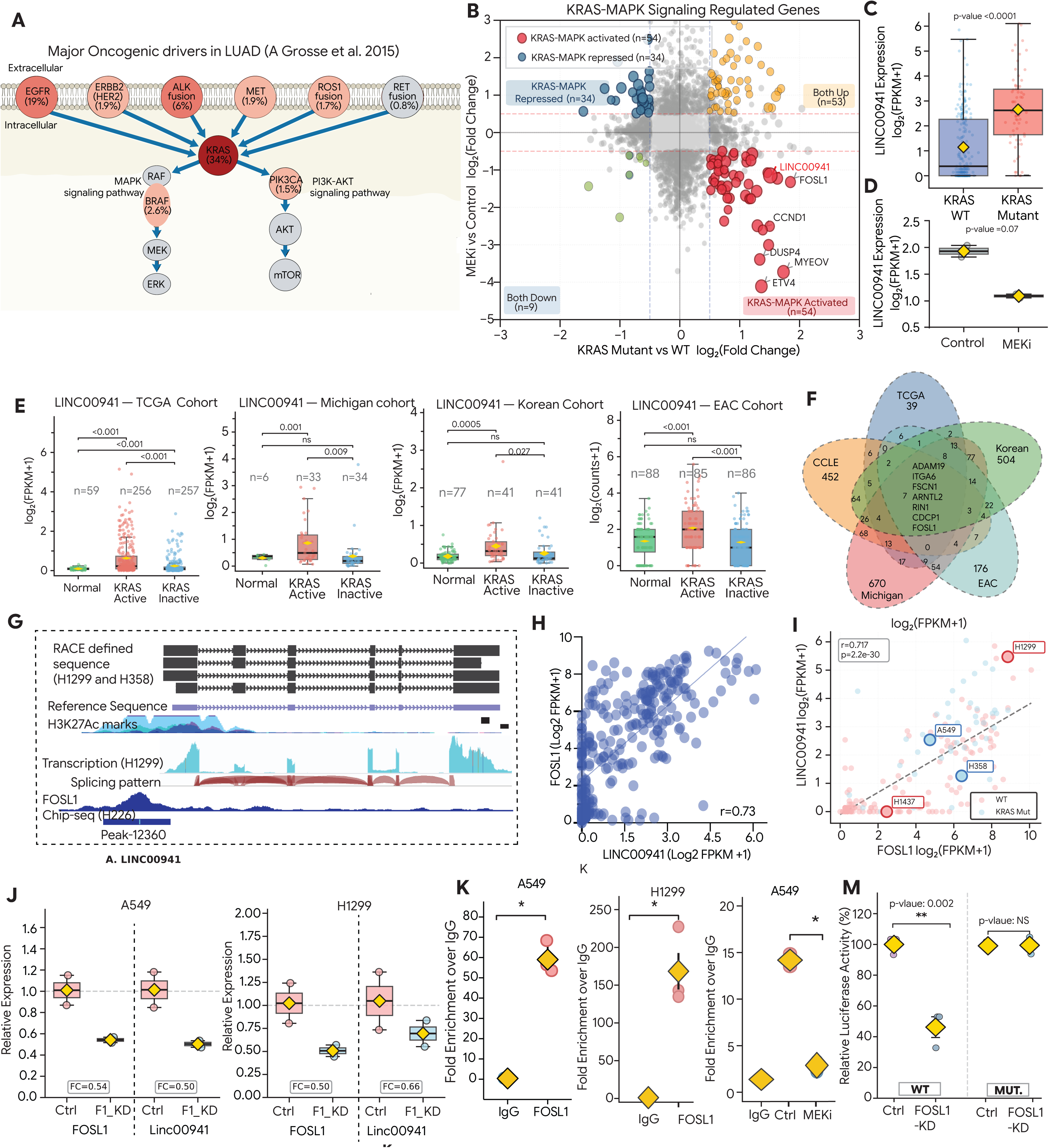
KRAS–MAPK signalling regulates long non-coding RNA LINC00941 expression in lung adenocarcinoma. **(A)** Mutation frequency of common driver genes (Grosse et al ) in lung adenocarcinoma cohorts, highlighting the prevalence of *KRAS* alterations. **(B)** Starburst plot identifying genes dysregulated by KRAS–MAPK signalling that are reversed upon MEKi treatment. Fold changes of differentially expressed genes identified from RNAseq data between MEKi vs vehicle control treated A549 cells (y-axis), compared to DEGs identified between KRAS-mutant and KRAS wild-type lung adenocarcinoma samples bulk RNAseq (x-axis). LINC00941 emerged as one of the top upregulated in KRAS mutant tumors, that was downregulated upon MEK inhibition in KRAS-mutant cells. **(C)** Box plot showing expression of LINC00941 in *KRAS* wild-type versus *KRAS*-mutant lung adenocarcinoma samples. **(D)** Expression of LINC00941 in control and MEK inhibitor–treated cells plotted as log_2_(FPKM + 1). **(E)** LINC00941 expression in normal lung tissues, *KRAS*-active, and *KRAS*-inactive patient samples across four independent lung adenocarcinoma cohorts. Patients were stratified into *KRAS*- active and *KRAS*-inactive groups using single-sample Gene Set Enrichment Analysis (ssGSEA) scores, validated by PROGENy-derived MAPK activity scores. **(F)** Venn diagram showing genes whose expression correlated with LINC00941 across four patient cohorts and a CCLE cell line cohort, identifying seven commonly correlated genes. **(G)** Genomic locus of LINC00941 visualised using the UCSC Genome Browser, showing H3K27ac enrichment at the promoter region in H1299 and H358 cells and a FOSL1 binding site identified by ChIP-seq analysis in H226 cells. Rapid amplification of cDNA Ends (RACE) defined transcripts are show as a track on top. **(H)** Correlation analysis of LINC00941 and FOSL1 expression in TCGA lung adenocarcinoma patient samples RNASeq Data. **(I)** Scatter plot showing correlation between LINC00941 and FOSL1 RNA expression across lung cancer cell lines. *KRAS*-mutant cell lines are indicated in blue and *KRAS* wild-type cell lines in red. **(J)** Relative expression of FOSL1 and LINC00941 following FOSL1 knockdown in A549 and H1299 cells. **(K)** ChIP-qPCR analysis showing enrichment of FOSL1 binding at the LINC00941 promoter in A549 and H1299 cells compared with IgG control. **(L)** ChIP-qPCR analysis of FOSL1 occupancy at the LINC00941 promoter in control and MEK inhibitor–treated H1299 cells. **(M)** Luciferase reporter assay showing activity of the LINC00941 promoter cloned upstream of luciferase in control and FOSL1-knockdown cells. Mutation of the FOSL1 binding site abolishes the effect of FOSL1 knockdown on promoter activity. Data are presented as mean ± SEM. Statistical significance was determined using Student’s *t*-test unless otherwise indicated. ^∗^*P <* 0.05; ^∗∗^*P <* 0.01; ^∗∗∗^*P <* 0.001.

To assess clinical relevance, we stratified LUAD patient samples from publicly available datasets into KRAS-active and KRAS-inactive groups using single-sample Gene Set Enrichment Analysis (ssGSEA) scores, validated by PROGENy-derived MAPK activity scores. LINC00941 expression was significantly higher in the KRAS-active group compared to the KRAS-inactive group (Figure 1E; *p <* 0.001). Consistent with this, LINC00941 expression was significantly elevated in LUAD patient samples relative to normal lung tissue across four independent patient cohorts (Figure 1E; all *p <* 0.05).

To investigate the transcriptional mechanism underlying LINC00941 upregulation, we performed Pearson correlation analysis between LINC00941 expression and all annotated genes across four patient datasets and one cell line dataset (CCLE). This unbiased approach identified seven genes consistently correlated with LINC00941 expression across all five datasets (Figure 1F). Among these, FOSL1—an AP-1 transcription factor and established downstream effector of KRAS-MAPK signalling (Supplementary Figure- 1B)—emerged as a top candidate. Examination of publicly available FOSL1 ChIP-seq data from the ENCODE dataset (accessed via the UCSC Genome Browser) revealed a prominent FOSL1 binding peak at the LINC00941 promoter (Figure 1G). Consistent with this, LINC00941 expression correlated strongly with FOSL1 expression (Pearson *r* = 0.73) across LUAD patient samples (Figure 1H). Analysis of NSCLC cell lines further confirmed this relationship: KRAS-wild-type cell lines showed lower LINC00941 expression, while KRAS-mutant lines with elevated FOSL1 expression exhibited correspondingly high LINC00941 levels (Figure 1I).

To establish a direct regulatory relationship, we performed FOSL1 knockdown in A549 and H1299 cells and observed a significant reduction in LINC00941 expression (Figure 1J; *p <* 0.05), confirming that FOSL1 is required for LINC00941 transcription. Overexpression of the MEK– ERK2 fusion significantly upregulated both FOSL1 and LINC00941 expression relative to empty-vector controls (Supplementary Figure 1C and D), demonstrating that activation of the MEK–ERK axis is sufficient to drive LINC00941 transcription through FOSL1. To directly test whether ERK activity is the critical downstream node, cells overexpressing the MEK–ERK2 fusion were subsequently treated with a MEK inhibitor. Strikingly, MEK inhibitor treatment failed to suppress LINC00941 expression in the presence of the MEK–ERK2 fusion, resulting in significant rescue of LINC00941 levels compared to inhibitor-treated empty-vector controls (Supplementary Figure 1E). To determine whether FOSL1 directly occupies the LINC00941 promoter, we performed chromatin immunoprecipitation (ChIP) assays in A549 and H1299 cells. FOSL1 ChIP yielded greater than 50-fold enrichment of the LINC00941 promoter region relative to IgG control in both cell lines (Figure 1K). MEK inhibition strongly reduced FOSL1 binding at the LINC00941 promoter compared to vehicle-treated controls (Figure 1L), demonstrating that FOSL1 recruitment is MEK-dependent.

To functionally validate FOSL1-mediated transcriptional control of LINC00941, we cloned the LINC00941 promoter upstream of a luciferase reporter in the pGL4 basic vector. FOSL1 knockdown resulted in a significant decrease in luciferase activity driven by the wild-type promoter (Figure 1M; *p <* 0.05). Importantly, mutation of the FOSL1 binding site abolished this response (Figure 1M). Collectively, these data establish that LINC00941 is a direct transcriptional target of FOSL1, regulated by the KRAS-MAPK signalling pathway in LUAD. To determine whether FOSL1-dependent regulation of LINC00941 extends beyond *KRAS*-mutant contexts, we confirmed FOSL1 occupancy of the LINC00941 promoter and FOSL1-dependent transcription in H1299 cells harbouring an NRAS^Q61K^ mutation, demonstrating that LINC00941 regulation is a shared consequence of oncogenic RAS–MAPK–FOSL1 signalling rather than a *KRAS* allele-specific phenomenon.

### LINC00941 is required for lung adenocarcinoma cell growth, and its depletion induces stress-induced premature senescence

To determine the functional requirement of LINC00941 in LUAD, we first validated its expression across a panel of NSCLC cell lines and selected three lines with high endogenous LINC00941 levels—H358 and A549 (both KRAS-mutant) and H1299 (harboring an NRAS^Q61K^ mutation, which equivalently activates the RAF–MEK–ERK cascade and induces comparable FOSL1 upregulation)— and one line with negligible expression (H1437) for functional studies (Figure 1I). The inclusion of both KRAS- and NRAS-mutant backgrounds was adopted deliberately to assess whether LINC00941 regulation and function represent a general consequence of oncogenic RAS–MAPK– FOSL1 hyperactivation rather than a KRAS allele-specific phenomenon. siRNA-mediated knockdown was performed using four independent siRNAs, and knockdown efficiency at the RNA level was confirmed for all constructs (Figure 2A). Relative to cells transfected with a non-targeting control siRNA (siNT), depletion of LINC00941 in H358, A549, and H1299 cells produced a significant reduction in cell proliferation and a marked suppression of colony formation (Figures 2B and 2C). Critically, neither phenotype was observed in the low-expressing H1437 line (Figure 2B), establishing that the antiproliferative effects are on-target and dependent on endogenous LINC00941 expression.

**Figure 2.**
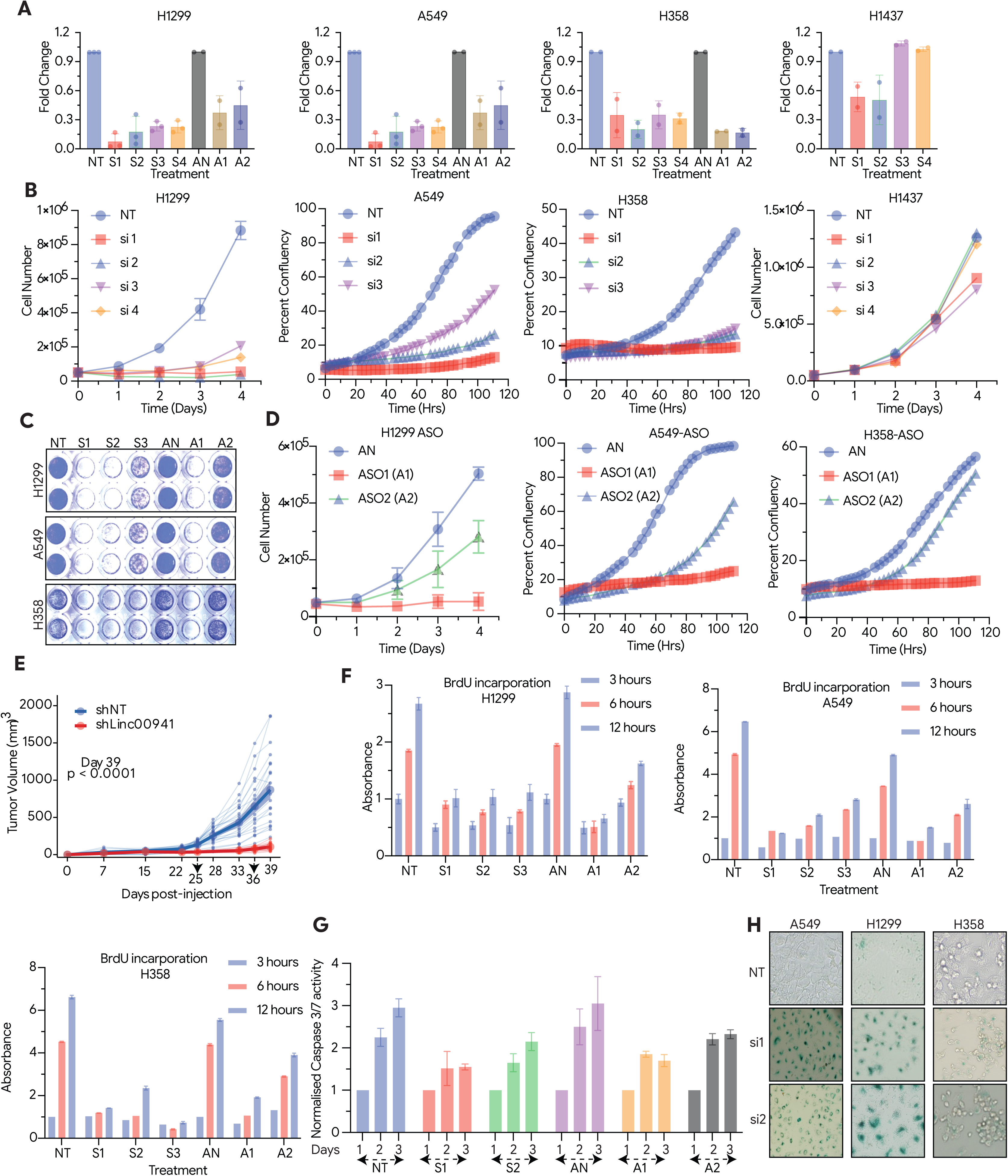
LINC00941 is required for proliferation and induces senescence in lung adenocarcinoma cells. **(A)** Quantitative RT-PCR analysis of LINC00941 expression in A549, H358, H1299, and H1437 cells following transfection with non-targeting siRNA (NT), non-targeting antisense oligonucleotide (AN), siLINC00941 (S1, S2,S3 and S4), or ASO-LINC00941 (A1, A2), confirming efficient knockdown. **(B)** Cell proliferation assays in A549, H358, H1299, and H1437 cells following transfection with siNT or siLINC00941. Proliferation was significantly reduced in A549, H358, and H1299 cells, whereas H1437 cells, which lack LINC00941 expression, showed no significant change. **(C)** Long-term colony formation assay in A549, H358, and H1299 cells treated with siNT, siL- INC00941, ASO-NT, or ASO-LINC00941. Cells were fixed and stained with crystal violet after two weeks. Representative images and quantification show a significant reduction in colony formation upon LINC00941 knockdown. **(D)** Cell proliferation analysis in A549, H358, and H1299 cells treated with ASO-NT or ASO- LINC00941, demonstrating a significant decrease in proliferation following LINC00941 depletion. **(E)** *In vivo* tumour growth assay in which H1299 cells stably expressing control shRNA or shLINC00941 were injected subcutaneously into NOD/SCID mice. Tumour volume was measured at the indicated time points, showing significantly reduced tumour growth upon LINC00941 knockdown. **(F)** BrdU incorporation assay in A549, H358, and H1299 cells following treatment with siNT, siL- INC00941, ASO-NT, or ASO-LINC00941, indicating reduced DNA synthesis upon LINC00941 depletion. **(G)** Caspase-3/7 activity measured using a Promega luminescence assay in H1299 cells treated with siNT, siLINC00941, ASO-NT, or ASO-LINC00941. No significant increase in apoptotic activity was observed following LINC00941 knockdown. **(H)** Senescence-associated *β*-galactosidase staining in A549, H1299, and H358 cells transfected with siNT or siLINC00941. LINC00941 depletion resulted in a significant increase in *β*-galactosidase-positive cells compared with controls. Data are presented as mean ± SEM. Statistical significance was determined using Student’s *t*-test unless otherwise indicated. ^∗^*P <* 0.05; ^∗∗^*P <* 0.01; ^∗∗∗^*P <* 0.001.

To exclude siRNA off-target effects and orthogonally validate these findings, we employed two independent antisense oligonucleotides (ASOs) alongside a non-targeting ASO control (AN), with knockdown efficiency confirmed (Figure 2A). ASO-mediated depletion of LINC00941 recapitulated the siRNA phenotype, producing a significant decrease in cell proliferation and colony numbers across all three expressing cell lines (Figures 2C and 2D). To further validate the growth-suppressive phenotype using a stable genetic system, shRNA-mediated knockdown of LINC00941 was confirmed in both H1299 and A549 cells and similarly reduced cell proliferation with ASO1 showing stronger effect than ASO2 *in vitro* (Supplementary Figure 2 A–C)

To establish the requirement of LINC00941 for tumour growth *in vivo*, H1299 cells stably expressing shLINC00941 or a non-targeting shRNA control (shNT) were implanted subcutaneously into NOD-SCID mice (shNT, *n* = 20 tumours; shLINC00941, *n* = 17 tumours). Tumour volumes were monitored over 39 days. shNT-implanted animals developed progressively growing tumours, reaching a mean volume of 870.2 ± 519.4 mm^3^ at the experimental endpoint. In contrast, shLINC00941 tumours exhibited markedly impaired growth, with a mean endpoint volume of 104.3 ± 47.8 mm^3^ — representing an 88% reduction and an 8.35-fold difference relative to controls (*p* = 3.7 × 10^−7^; Figure 2E). Significant divergence in growth trajectories was apparent from Day 22 onwards and was sustained at all subsequent measurement timepoints (*p <* 0.0001), demonstrating that LINC00941 expression is required for tumour establishment and progression *in vivo*.

To characterise the mechanistic basis of growth suppression, we performed complementary cell cycle and viability analyses. A 5-bromo-2^′^-deoxyuridine (BrdU) incorporation assay revealed a significant decrease in active DNA synthesis following LINC00941 knockdown in H358, A549, and H1299 cells (Figure 2F; *p <* 0.001), indicating impaired replication fork activity. Consistent with this, flow cytometry-based cell cycle profiling by propidium iodide (PI) staining demonstrated a significant accumulation of cells in the S and G_2_/M fractions with a concomitant reduction in the G_1_ population in both siRNA- and ASO-treated cells relative to matched controls (Supplementary Figure 2D; *p <* 0.05). The concurrent reduction in BrdU incorporation indicates that the apparent S-phase accumulation reflects cells harbouring stalled or collapsed replication forks rather than active DNA synthesis — a profile consistent with replication stress and DNA damage, characterised further in subsequent sections. Notably, caspase-3/7 activity was not significantly elevated across two independent siRNAs and two ASOs compared to matched controls (Figure 2G), excluding apoptosis as a determinant of the observed growth inhibition. Consistent with a senescence response, LINC00941-depleted cells exhibited a significant increase in senescence-associated *β*- galactosidase (SA-*β*-gal) activity relative to control cells (Figure 2H). Collectively, the combination of impaired DNA replication, replication stress-associated cell cycle arrest, absence of apoptosis, and induction of cellular senescence establishes that LINC00941 loss triggers stress-induced premature senescence (SIPS) in LUAD cells.

### LINC00941 depletion suppresses DNA replication gene expression and induces senescence

Prior to investigating the functional role of LINC00941, we characterized the global transcriptional consequences of LINC00941 loss by performing RNA sequencing of H1299 cells following siRNA-mediated knockdown relative to a non-targeting control. Knockdown efficiency was confirmed at the transcriptome level, with LINC00941 itself among the most significantly depleted transcripts (log_2_ FC = −4.59; *p*_adj_ = 0.012; Figure 3A, B and supplementary table ST7 ). Of 15,208 expressed genes, differential expression analysis identified 586 significantly dysregulated genes (*p*_adj_ *<* 0.05; | log_2_ FC| *>* 1), comprising 503 downregulated and 83 upregulated genes (Figure 3A, B). To ensure robustness of the transcriptomic findings and exclude platform- or reagent-specific artefacts, RNA-seq-derived differentially expressed genes were independently validated across two microarray experiments using distinct siRNA sequences hybridised against non-targeting control (Figure 3A). The convergence of gene expression changes across three independent experiments — spanning two orthogonal platforms and two independent siRNAs — provides cross-platform, cross-reagent confirmation of the core transcriptional signature and substantially strengthens confidence in the identified gene sets. Pathway-level conclusions are supported by this multi-platform agreement, while individual gene-level findings from the RNA-seq dataset are further corroborated by the microarray data (Figure 3A).

**Figure 3.**
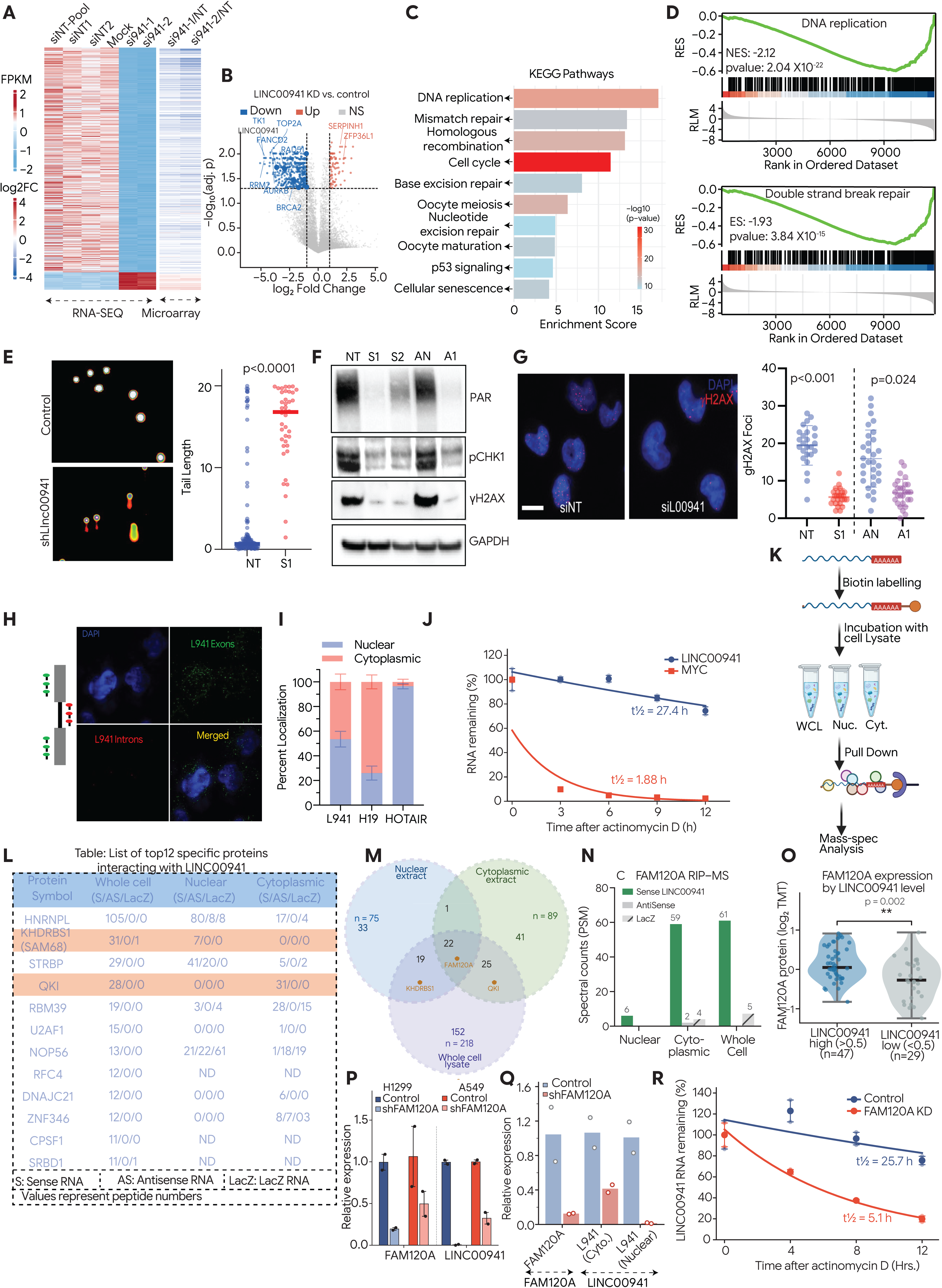
LINC00941 depletion alters cell cycle progression and DNA damage response in lung carcinoma cells. **(A)** Heat map of differentially expressed genes identified by RNA-seq analysis comparing LINC00941 knockdown and control cells. Heat map showing expression of the same set of differentially expressed genes obtained from microarray analysis comparing siLINC00941 and siNT conditions using two independent siRNAs. **(B)** Volcano plot showing differentially expressed genes following LINC00941 knockdown in H1299 cells. Cells were transfected with two independent siRNAs targeting LINC00941 or four non-targeting siRNAs (siNT) for 48 h prior to RNA isolation and RNA sequencing. **(c)** KEGG pathway enrichment analysis of differentially expressed genes upon LINC00941 knock-down. Bar height indicates enrichment score and colour represents − log_10_(*P* value). **(D)** Pre-ranked gene set enrichment analysis (GSEA) using Hallmark gene sets based on RNA-seq data. DNA replication and double-strand break repair pathways were among the most significantly negatively enriched gene sets following LINC00941 depletion. **(E)** Comet assay performed in control and LINC00941-depleted H1299 cells 48 h post-transfection. Representative images are shown and tail length was quantified using Comet Analyser software. **(F)** Immunoblot analysis of DNA damage response signalling proteins in H1299 cells treated with siNT, siLINC00941, ASO-NT, or ASO-LINC00941, demonstrating reduced activation of DDR pathways upon LINC00941 knockdown. **(G)** Immunofluorescence staining of *γ*H2AX in control and siLINC00941-treated H1299 cells. Representative images are shown and *γ*H2AX foci were quantified, revealing a significant reduction in foci number following LINC00941 depletion. **(H)** RNA fluorescence *in situ* hybridisation (RNA-FISH) analysis of LINC00941 localisation in H1299 cells. Probes targeting LINC00941 exons were labelled with Cy3 (green), and intronic probes were labelled with Cy5 (red). Nuclei were counterstained with DAPI. LINC00941 signal was detected in both the nuclear and cytoplasmic compartments as evident from the merged image. **(I)** Subcellular fractionation of H1299 cells followed by quantitative RT-PCR analysis of LINC00941 expression in nuclear and cytoplasmic fractions. H19 and HOTAIR were used as control lncRNAs for cytoplasmic and nuclear localisation, respectively. **(J)** RNA stability assay in H1299 cells treated with the transcriptional inhibitor actinomycin D. RNA was isolated at the indicated time points and LINC00941 expression was measured by quantitative RT-PCR and normalised to actin mRNA. **(K)** Schematic overview of the experimental workflow used to identify LINC00941-interacting proteins in whole-cell, nuclear, and cytoplasmic fractions. *In vitro*-transcribed, biotin-labelled LINC00941 RNA was incubated with cell lysates, followed by streptavidin pulldown and mass spectrometry analysis. **(L)** Table listing proteins enriched in LINC00941 pulldown samples from whole-cell, nuclear, and cytoplasmic lysates as identified by mass spectrometry. **(M)** Venn diagram showing overlap of LINC00941-associated proteins across nuclear and cyto-plasmic fractions. KHDRBS1 (SAM68) was uniquely enriched in the nuclear fraction, whereas QKI was enriched in the cytoplasmic fraction. **(N)** Bar graph depicting spectral counts from RNA immunoprecipitation (RIP), showing FAM120A binding to LINC00941 and [Control RNA] across nuclear, cytoplasmic, and whole-cell fractions. FAM120A demonstrated robust association with LINC00941 in all three fractions, while binding to [Control RNA] was negligible, confirming the specificity of the FAM120A–LINC00941 interaction. **(O)** Violin plot illustrating the differential expression of FAM120A protein in patient samples stratified by LINC00941 expression levels (high versus low). FAM120A expression was significantly elevated in LINC00941-high samples compared to LINC00941-low samples, indicating a positive correlation between FAM120A abundance and LINC00941 expression in clinical specimens. **(P)** Bar graph showing LINC00941 RNA and FAM120A RNA levels upon FAM120A knockdown relative to control cells. FAM120A depletion resulted in a significant reduction in LINC00941 RNA abundance, suggesting that FAM120A positively regulates LINC00941 expression or stability. **(Q)** RT-qPCR analysis of LINC00941 RNA expression in nuclear and cytoplasmic fractions of H1299 cells following FAM120A knockdown. RNA levels are presented relative to control. **(R)** Actinomycin D chase assay in control and FAM120A-depleted H1299 cells. Transcription was blocked with actinomycin D and LINC00941 RNA levels were measured by RT-qPCR at 0, 4, 8, and 12 hours. Values are normalised to the 0 h timepoint and expressed as percentage RNA remaining. Exponential decay fitting yielded half-lives of 25.7 h (control) and 5.1 h (FAM120A KD). Data are presented as mean ± SEM unless otherwise indicated. Statistical significance was determined using Student’s *t*-test. ^∗^*P <* 0.05; ^∗∗^*P <* 0.01; ^∗∗∗^*P <* 0.001.

Gene ontology and pathway classification of downregulated genes revealed a striking enrichment for processes governing DNA replication and cell cycle progression (Figure 3C). The most severely suppressed transcripts included rate-limiting enzymes of deoxyribonucleotide biosynthesis— *TK1* (log_2_ FC = −4.75; *p*_adj_ = 0.012), *RRM2* (log_2_ FC = −4.68; *p*_adj_ = 0.012), *TYMS* (log_2_ FC = −3.83; *p*_adj_ = 0.010), and *DHFR* (log_2_ FC = −3.44; *p*_adj_ = 0.039)—whose coordinated suppression is sufficient to restrict dNTP pools and impair DNA synthesis. Replication fork components were equally affected, including *CDC45* (log_2_ FC = −4.47), *CLSPN* (−3.95), *HJURP* (−3.92), *TOP2A* (−4.16), *AURKB* (−3.99), *MKI67* (−3.98), and *PCNA* (−2.48). In total, 38 genes with established roles in cell cycle regulation and DNA replication were significantly downregulated, spanning origin licensing (*ORC1*, *CDC6*, *CDT1*), the CMG helicase complex (*GINS1*, *GINS2*, *GINS4*, *CDC45*, *MCM6*), replication fork elongation (*POLD1*, *POLA1*, *POLA2*, *PRIM1*, *LIG1*), and mitotic progression (*CDK1*, *CEP55*, *AURKB*, *TTK*). Additionally, six genes mediating homologous recombination-based replication fork restart were significantly downregulated, including the Fanconi anaemia pathway components *FANCD2* (log_2_ FC = −3.63; *p*_adj_ = 0.019), *FANCI* (−3.34), and *FANCB* (−2.17), as well as the recombinase *RAD51* (−2.94; *p*_adj_ = 0.020) and *BRCA2* (−2.10; *p*_adj_ = 0.035).

Consistent with these findings, Gene Set Enrichment Analysis (GSEA) demonstrated significant negative enrichment for both the DNA replication gene set (NES = −2.12; *p* = 2.04 × 10^−22^; Figure 3D) and the DNA repair gene set (NES = −1.93; *p* = 3.84 × 10^−15^; Figure 3D) in siLINC00941 cells.

Given the broad transcriptional suppression of DNA repair genes, we investigated whether LINC00941 depletion functionally impairs the DNA damage response (DDR). Alkaline comet assays revealed a greater than ten-fold increase in comet tail length in LINC00941-depleted cells relative to non-targeting controls (*p <* 0.001; Figure 3E), demonstrating a marked accumulation of unrepaired DNA strand breaks. Western blot analysis revealed that knockdown of LINC00941 abrogated phosphorylation of CHK1 (pCHK1) and reduced steady-state levels of *γ*H2AX and poly-ADP-ribose (PAR) (Figure 3F), indicating impaired checkpoint kinase signalling and PARP-mediated single-strand break repair. Immunofluorescence analysis further demonstrated a significant reduction in *γ*H2AX foci (Figure 3G) and 53BP1 foci (Supplementary Figure 3A) in LINC00941-depleted cells. Collectively, these findings establish that LINC00941 is required for competent activation of the DDR.

Among the 83 upregulated genes, several are associated with cellular senescence and stress response pathways, including *SERPINH1* (log_2_ FC = +1.91; *p*_adj_ = 0.007), *CEBPD* (+1.87), *ZFP36L1* (+2.20), *HIST1H4H* (+2.82), *HIST1H2BK* (+2.08), and *NATD1* (+2.39; Figure 3A, B), consistent with the senescence programme identified by SA-*β*-galactosidase staining in LINC00941- depleted cells (Figure 2H).

### LINC00941 is an exceptionally stable and universally distributed transcript

To characterise the subcellular distribution of LINC00941, RNA fluorescence *in situ* hybridisation (RNA-FISH) was performed in H1299 cells. Signal was detected in both nuclear and cytoplasmic compartments at comparable intensity, indicating a dual subcellular localisation (Figure 3H). Quantitative subcellular fractionation followed by strand-specific RT-qPCR confirmed this near-equal partitioning across compartments (Figure 3I).

To assess intrinsic RNA stability, H1299 cells were treated with the transcription inhibitor actinomycin D and LINC00941 abundance was monitored at successive time points by RT-qPCR. LINC00941 exhibited exceptional stability, with an estimated half-life of approximately 27 hours (Figure 3J), a property that is uncommon among long non-coding RNAs [16, 17].

Given its stability and dual compartmentalisation, we reasoned that LINC00941 may exert distinct mechanistic functions dependent on its subcellular localisation and interacting protein partners. To resolve compartment-specific interactions, RNA pulldown coupled with quantitative mass spectrometry was performed on nuclear and cytoplasmic extracts as well as whole-cell lysate (WCL) from H1299 cells, using sense LINC00941 RNA as bait alongside antisense and LacZ control probes (Figure 3K). Candidate interactors were prioritised based on the specificity and magnitude of enrichment in the sense pulldown relative to both control probes, compartmental consistency, and established functional relevance to RNA biology and cancer (Supplementary Table ST8).

Three proteins satisfied these criteria. FAM120A, a nucleocytoplasmic scaffold protein implicated in mRNA stabilisation, was the strong interactor enriched across all three fractions — nuclear (6 spectral counts), cytoplasmic (59 counts), and WCL (61 counts) — with complete absence of signal in the antisense control, providing stringent evidence of binding specificity. KHDRBS1 (SAM68), a STAR-family KH-domain RNA-binding protein with established roles in DNA damage response and tumour progression, was identified as a nuclear-enriched interactor, yielding 7 and 31 spectral counts in the nuclear and WCL sense pulldowns, respectively, with no signal detected in either control or in the cytoplasmic fraction. QKI, a KH-domain RNA-binding protein that post-transcriptionally regulates splicing and mRNA localisation with context-dependent oncogenic and tumour-suppressive functions, was identified as a cytoplasm-specific interactor, with 30 spectral counts in the cytoplasmic sense pulldown against a background of zero in both controls, and confirmed in the WCL fraction (28 counts). These three proteins were selected for downstream functional validation (Figure 3L, M and Supplementary Figure 3B).

### FAM120A binds LINC00941 and regulates its RNA stability

Having demonstrated that LINC00941 exhibits exceptional RNA stability comparable to that of coding mRNAs — an uncommon property for long non-coding RNAs [16, 17] — we sought to investigate the molecular basis of this stability. Our subcellular RNA pulldown mass spectrometry identified FAM120A as a high-confidence LINC00941-interacting protein enriched in both nuclear and cytoplasmic fractions, and given its established role in protecting target RNAs from degradation [18, 19], we hypothesised that FAM120A may be responsible for maintaining LINC00941 stability in lung cancer cells.

To explore this possibility, the relationship between LINC00941 RNA expression and FAM120A protein levels was first examined across lung cancer patient samples. A significant positive correlation was observed: FAM120A protein levels were consistently elevated in samples with high LINC00941 expression relative to those with low expression (Figure 3N), suggestive of coordinated regulation between the two molecules.

To directly assess whether FAM120A is required for LINC00941 stability, FAM120A was depleted by RNA interference in H1299 and A549 lung cancer cells and LINC00941 RNA levels were subsequently quantified by RT-qPCR. FAM120A depletion produced a marked reduction in LINC00941 RNA abundance (Figure 3P). Knockdown of FAM120A resulted in a marked reduction of LINC00941 expression in both the cytoplasmic and nuclear fractions (Figure 3Q) . In contrast, knockdown of the other LINC00941-interacting proteins KHDRBS1 or QKI had no significant effect on LINC00941 levels, implicating FAM120A specifically in the post-transcriptional maintenance of LINC00941 (Supplementary figure 3C–E).To confirm this at the level of RNA turnover, actinomycin D chase experiments were performed in control and FAM120A-depleted H1299 cells. Exponential decay fitting revealed a LINC00941 half-life of approximately 25.7 hours in control cells, consistent with its previously characterised stability, whereas FAM120A knockdown reduced the half-life to 5.1 hours — a five-fold acceleration in RNA decay (Figure 3R). Collectively, these findings establish that FAM120A binds LINC00941 across subcellular compartments and acts as a critical stabilising factor that protects LINC00941 from degradation in lung cancer cells.

### LINC00941-SAM68 nuclear interaction, promotes SAM68 stability by rendering protection from proteasomal degradation

To validate the nuclear interaction, RNA pull-down assays using biotin-labelled sense LINC00941, antisense LINC00941, and LacZ RNA as specificity controls confirmed enrichment of SAM68 protein specifically in the sense LINC00941 fraction, with a corresponding depletion from the unbound supernatant, in both H1299 and A549 cells (Figure 4A). RNA immunoprecipitation (RIP) corroborated this, with anti-SAM68 antibody enriching LINC00941 transcripts *>*500-fold above IgG control (Figure 4B). To confirm that this interaction is spatially restricted to the nuclear compartment, we performed RIP using subcellular-fractionated lysates. SAM68 immunoprecipitation enriched LINC00941 11.1-fold above input in the nuclear fraction, compared with only 2.7- fold in the cytoplasmic fraction (Figure 4E). Reciprocally, QKI immunoprecipitation enriched LINC00941 4.4-fold in the cytoplasmic fraction versus 2.8-fold in the nuclear fraction (Figure 4C). This compartment-resolved RIP establishes that LINC00941 engages SAM68 predominantly in the nucleus and QKI predominantly in the cytoplasm.

**Figure 4.**
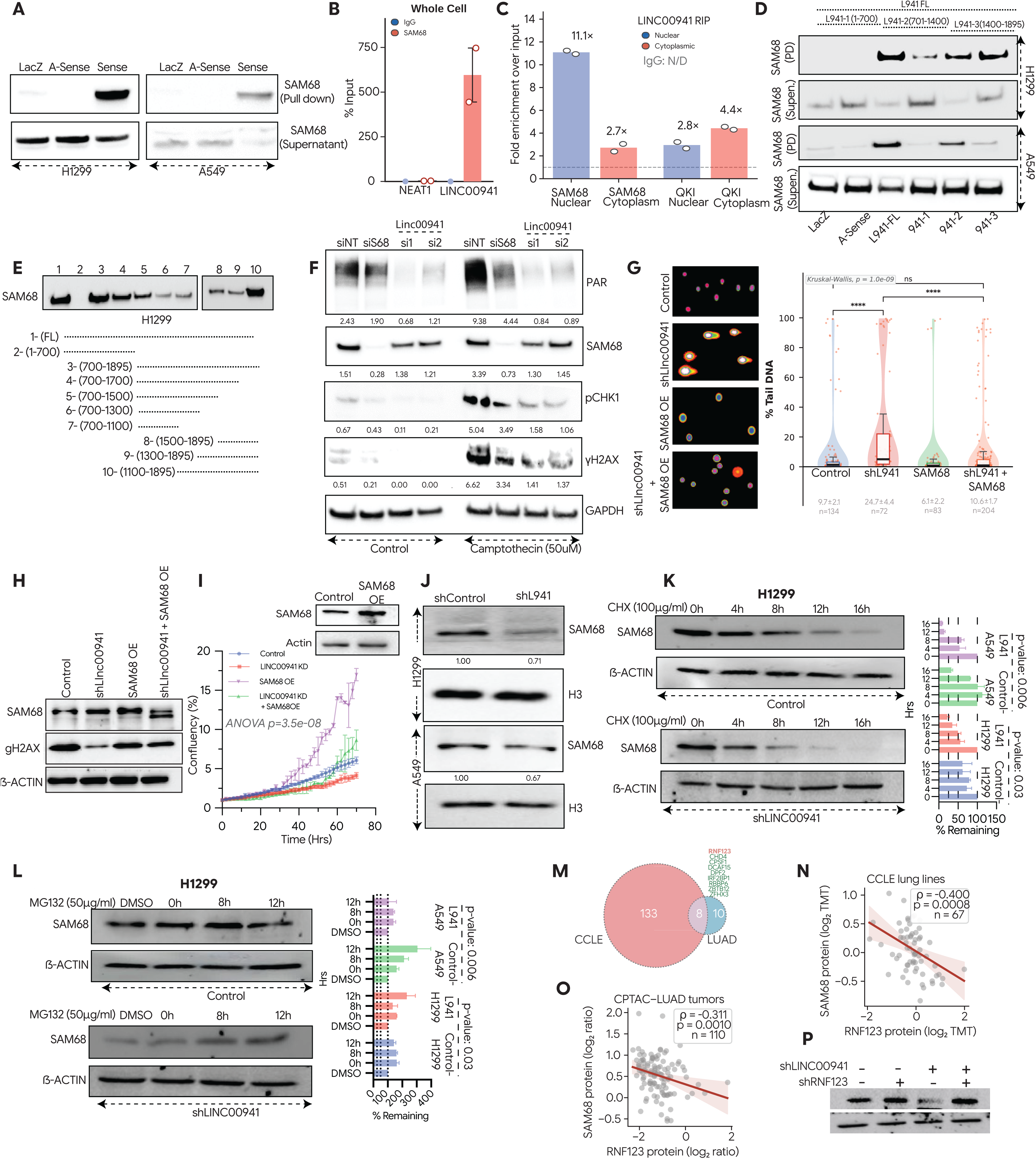
LINC00941 interacts with SAM68 to regulate the DNA damage response in lung cancer cells. **(A)** RNA pulldown followed by immunoblot analysis demonstrating interaction between biotin-labelled LINC00941 and SAM68 in H1299 and A549 cells. LacZ RNA and antisense LINC00941 RNA were used as negative controls. **(B)** Recipocal RNA immunoprecipitation (RIP) assay performed using anti-SAM68 antibody in H1299 cells to assess enrichment of LINC00941. IgG was used as a negative control. NEAT1 was included as negative controls for SAM68 binding. **(C)** RIP enrichment of LINC00941 by SAM68 and QKI in nuclear and cytoplasmic fractions. Fold enrichment = 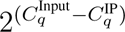. IgG controls showed no detectable amplification (not plotted). Dashed line = input level (1×). **(D)** RNA pulldown assays using three truncated LINC00941 constructs to map the SAM68 interaction domain. Loss of the 3^′^ region of LINC00941 markedly reduced SAM68 binding in both A549 and H1299 cells. LacZ RNA and antisense LINC00941 RNA were used as negative controls. PD: Pull down and Supern: Supernatant. **(E)** Fine mapping of the SAM68 binding region using ten overlapping LINC00941 RNA fragments. Pulldown analysis identified nucleotides 700–1800 as necessary for SAM68 interaction. **(F)** Immunoblot analysis of DNA damage response proteins in control, SAM68-depleted, and LINC00941-depleted cells treated with or without the DNA-damaging agent camptothecin. Blots were probed for PAR, SAM68, phospho-CHK1, *γ*H2AX, and GAPDH as a loading control. **(G)** Comet assay assessing DNA damage in control cells, shLINC00941 cells, SAM68-overexpressing cells, and shLINC00941 cells with SAM68 overexpression. Quantification shows significant rescue of DNA damage upon SAM68 overexpression in LINC00941-depleted cells. **(H)** Immunoblot analysis of *γ*H2AX levels in control, shLINC00941, SAM68-overexpressing, and shLINC00941 + SAM68-overexpressing cells, confirming reduced DNA damage signalling upon SAM68 rescue. **(I)** Cell proliferation assay performed in control cells, LINC00941-knockdown cells, SAM68-overexpressing cells, and cells with combined LINC00941 knockdown and SAM68 overexpression, demonstrating phenotypic rescue by SAM68. **(J)** SAM68 expression was measured in control and LINC00941-knockdown H1299 using western blot. **(K)** Cycloheximide chase assay assessing SAM68 protein stability in control and shLINC00941 H1299 cells. Cells were harvested at the indicated time points following cycloheximide treatment and SAM68 protein levels were analysed by immunoblotting. **(L)** Proteasome inhibition assay showing stabilisation of SAM68 protein in control and shLINC00941 cells treated with MG132, indicating proteasome-mediated degradation of SAM68 upon LINC00941 depletion. **(M)** Venn diagram showing the overlap of E3 ubiquitin ligases negatively correlated (R < 0.3) with SAM68 protein expression in Cancer Cell Line Encyclopedia (CCLE) cell lines (n = 54 E3 ligases) and Clinical Proteomic Tumor Analysis Consortium (CPTAC) lung adenocarcinoma (LUAD) tumors (n = 2 E3 ligases). RNF123 is the only E3 ligase common to both datasets. **(N)** Scatter plot showing negative correlation between SAM68 and RNF123 protein expression levels in CCLE cell lines. Pearson correlation coefficient (R) and p-value are indicated. **(O)** Scatter plot showing negative correlation between SAM68 and RNF123 protein expression levels in CPTAC LUAD tumor samples. Pearson correlation coefficient (R) and p-value are indicated. **(P)** Western blot analysis of SAM68 protein levels in NSCLC cells following RNF123 knockdown (shRNF123), LINC00941 knockdown (shLINC00941), or co-depletion of both RNF123 and LINC00941 (shRNF123 + shLINC00941). RNF123 knockdown alone does not alter SAM68 levels, whereas LINC00941 depletion reduces SAM68 protein expression. Co-depletion of RNF123 rescues SAM68 protein levels in LINC00941-knockdown cells. *β*-actin serves as loading control. Data are presented as mean ± SEM unless otherwise indicated. Statistical significance was determined using Student’s *t*-test. ^∗^*P <* 0.05; ^∗∗^*P <* 0.01; ^∗∗∗^*P <* 0.001.

To delineate the region of LINC00941 responsible for SAM68 interaction, we performed pulldown assays using truncated LINC00941 constructs. The N-terminal fragment spanning nucleotides 1–700 failed to pull down SAM68, whereas fragments spanning nucleotides 701–1400 and 1400– 1895 both retained binding capacity. Fine-mapping experiments using ten overlapping deletion constructs in H1299 cells identified a primary binding domain (BD-1) within nucleotides 700–1100 and a second discrete binding domain (BD-2) within the 3^′^ terminal region (nucleotides 1100– 1895; Figure 4D, E). The stronger SAM68 pulldown observed with full-length transcript compared to individual fragments suggests that BD-1 and BD-2 act cooperatively to facilitate high-affinity interaction.

Knockdown of SAM68 phenocopied the DDR defects observed upon LINC00941 depletion, including reduced *γ*H2AX foci and impaired CHK1 phosphorylation and cell proliferation (Figure 4F and Supplementary Figure 3F), implicating SAM68 as a downstream effector of LINC00941 in the DDR [20]. Critically, overexpression of SAM68 in LINC00941-depleted cells significantly rescued DNA strand break accumulation (comet tail moment: 24.7 ± 4.4 in siLINC00941 vs. 10.6 ± 1.7 in siLINC00941 + SAM68-OE vs. 9.7 ± 2.1 in siNT control; Figure 4G), and restored *γ*H2AX phosphorylation (Figure 4H) and cell proliferation (Figure 4I), establishing SAM68 as a critical downstream effector through which LINC00941 sustains tumour cell proliferation.

Knockdonw of LINC00941 resulted in a marked reduction in SAM68 protein (*>*30% decrease by densitometry; Figure 4F J,Supplementary Figure 3G) without a corresponding change in *KHDRBS1* mRNA abundance (Supplementary Figure 3H), suggesting post-transcriptional regulation. Cycloheximide chase assays demonstrated that LINC00941 depletion reduced the SAM68 half-life from *>*8 hours to ∼4 hours in both H1299 and A549 cells (Figure 4K and Supplementary figure 3I), indicating that LINC00941 substantially protects SAM68 from proteolytic turnover. MG132 treatment fully rescued SAM68 protein levels in LINC00941-depleted cells (Figure 4L and Supplementary figure 3J), establishing that LINC00941 shields SAM68 from proteasome-mediated degradation. To identify the E3 ubiquitin ligase involved in SAM68 protein regulation, we performed correlation analysis between SAM68 protein expression and the expression of all annotated E3 ligases using two independent datasets: Cancer Cell Line Encyclopedia (CCLE) cell lines and the Clinical Proteomic Tumor Analysis Consortium (CPTAC) lung adenocarcinoma (LUAD) dataset. Using a stringent negative correlation threshold of R < 0.3, we identified 54 E3 ligases in CCLE cell lines and two E3 ligases in CPTAC LUAD tumors that negatively correlated with SAM68 protein levels. Only one E3 ligase, RNF123, was common to both datasets (Figure 4M), suggesting its potential involvement in SAM68 degradation. Scatter plots depicting the negative correlation between SAM68 and RNF123 protein expression in CCLE cell lines and CPTAC LUAD tumors are shown in Figures 4N and 4O, respectively. To functionally validate whether RNF123 regulates SAM68 protein stability in a LINC00941-dependent manner, we depleted RNF123 using short hairpin RNA (shRNA) and assessed SAM68 protein levels by Western blotting. In control conditions, RNF123 knockdown alone did not significantly alter SAM68 protein expression (Figure 4P). As expected, depletion of LINC00941 resulted in a significant reduction in SAM68 protein levels. Importantly, co-depletion of RNF123 in LINC00941-knockdown cells rescued SAM68 protein expression, restoring it to near-control levels (Figure 4P). These results demonstrate that RNF123 mediates the degradation of SAM68 at the protein level, and that LINC00941 stabilizes SAM68 by preventing RNF123-mediated proteasomal degradation.

### LINC00941 sequesters QKI in the cytoplasm to prevent aberrant nuclear splicing activity

Our compartment-specific RNA pull-down screen identified the RNA-binding protein Quaking (QKI) as the primary cytoplasmic interaction partner of LINC00941. RNA pull-down assays using biotin-labeled sense LINC00941 confirmed specific enrichment of QKI protein relative to antisense LINC00941 and LacZ controls (SNF5 was used as a negative control) (Figure 5A). Bioinformatic analysis of the LINC00941 transcript identified two putative QKI response elements (QREs; 5^′^- NACUAAY-N_1−20_-UAAY-3^′^) located within the 3^′^ region. Pull-down assays using LINC00941 deletion constructs confirmed that a construct spanning nucleotides 810–1890 retained strong QKI binding, whereas a construct comprising nucleotides 1–810 failed to interact with QKI (Figure 5B). Reciprocal RIP using an anti-QKI antibody enriched LINC00941 transcripts 300-fold above IgG control, comparable to the canonical QKI target *NUMB* and substantially above non-target controls *THOR* and *Linc00857* (Figure 5C). Reciprocal RIPs using HUR and SNRNP70 antibodies pull downs served as additional controls, and the data collectively show very selective interaction between QKI protein and LINC00941 as compared to Other RNAs like Linc00857 and THOR (NUMB was used as a positive control for QKI binding), confirming the specificity of the LINC00941–QKI interaction.

**Figure 5.**
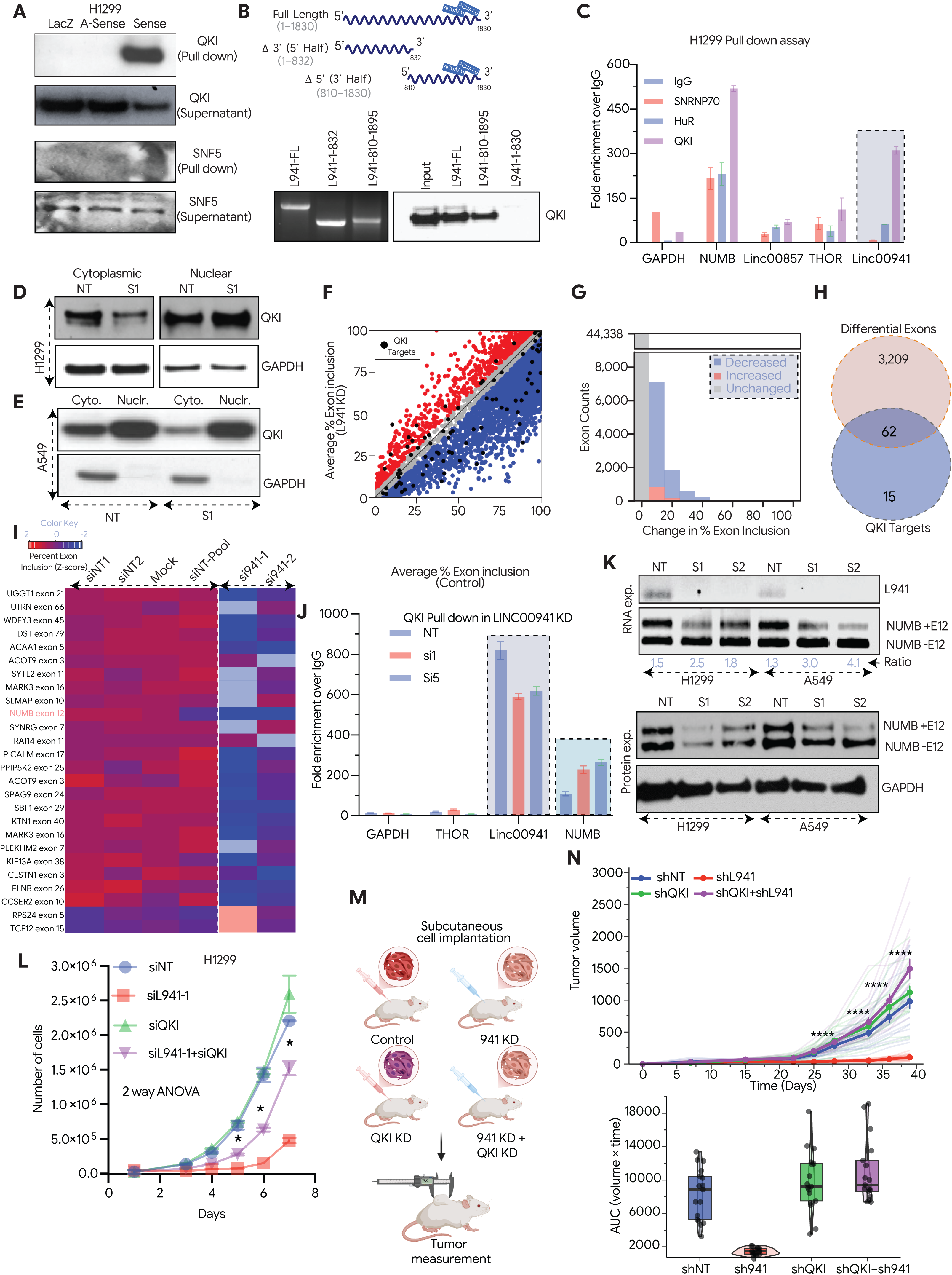
LINC00941 interacts with QKI in the cytoplasm and inhibits its nuclear localisation to regulate alternative splicing. **(A)** RNA pulldown assay demonstrating interaction between LINC00941 and QKI. *In vitro*-transcribed, biotin-labelled LINC00941 RNA was incubated with cell lysates, followed by streptavidin pulldown and immunoblotting for QKI. SNF5 was used as a negative control. **(B)** Mapping of the QKI-binding region on LINC00941 using full-length, 5^′^-half, and 3^′^-half RNA constructs. RNA pulldown followed by immunoblotting revealed that the 3^′^ half of LINC00941, which contains two predicted QKI-binding sites, is required for QKI interaction. **(C)** Recipocal RNA immunoprecipitation (RIP) assay performed using anti-QKI antibody in H1299 cells to assess enrichment of LINC00941. IgG was used as a negative control. SNRNP70 and HUR served as additional negative controls, whereas *NUMB* RNA served as a positive control. GAPDH, LINC00857, and THOR were included as negative controls for QKI binding. **(D)** Subcellular fractionation of control and siLINC00941-treated H1299 cells followed by immunoblotting for QKI. Loss of LINC00941 resulted in increased nuclear localisation of QKI. **(E)** Subcellular fractionation of control and siLINC00941-treated A549 cells followed by immunoblotting for QKI. Loss of LINC00941 resulted in increased nuclear localisation of QKI. **(F)** Alternative splicing analysis of RNA-seq data following LINC00941 knockdown. Differential exon inclusion and exclusion events were identified, with QKI target genes highlighted in black. **(G)** Quantification of splicing changes showing exon counts plotted against changes in percentage exon inclusion. Blue indicates decreased exon inclusion and red indicates increased exon inclusion. **(H)** Overlap between QKI target genes and differentially spliced exons identified upon LINC00941 depletion. Twenty-four of seventy-five QKI targets showed significant enrichment. **(I)** Heat map showing *Z*-score-normalised percentage exon inclusion for top QKI target genes. *NUMB* was identified as one of the most differentially spliced transcripts following LINC00941 knockdown. **(J)** QKI pulldown assay performed in control and LINC00941-knockdown cells to assess binding of QKI to *NUMB* RNA. GAPDH and THOR were used as negative controls. Increased QKI binding to *NUMB* was observed upon LINC00941 depletion. **(K)** RT-PCR analysis of *NUMB* splice variants with and without exon 12 in H1299 and A549 cells. LINC00941 knockdown significantly increased the exon-12-skipped isoform, which was further confirmed by immunoblotting showing increased protein levels of the corresponding NUMB isoform. **(L)** Rescue experiment assessing cell proliferation following combined knockdown of LINC00941 and QKI. Simultaneous depletion of QKI significantly rescued the proliferation defect induced by LINC00941 knockdown. **(M)** *In vivo* Schematics of rescue experiment in which control and LINC00941-knockdown cells, with or without QKI knockdown, were injected into immunocompromised mice. **(N)** *In vivo* Control and LINC00941-knockdown cells, with or without QKI knockdown, were injected into immunocompromised mice and Tumour volume was measured over time and area-under-the-curve (AUC) analysis demonstrated significant rescue of tumour growth upon combined LINC00941 and QKI depletion. **The shcontrol and shLINC00941 data is taken from** figure 2E**, both the data are part of one experiment** Data are presented as mean ± SEM unless otherwise indicated. Statistical significance was determined using Student’s *t*-test. ^∗^*P <* 0.05; ^∗∗^*P <* 0.01; ^∗∗∗^*P <* 0.001.

QKI shuttles between the cytoplasm and nucleus, and its nuclear accumulation drives alternative splicing of target pre-mRNAs. We hypothesised that cytoplasmic LINC00941 retains QKI in this compartment. Knockdown of LINC00941 resulted in 1.66-fold (H1299) and 6.2-fold (A549) increases in nuclear QKI with a corresponding reduction in cytoplasmic QKI (Figure 5D, E), demonstrating that LINC00941 is required for cytoplasmic retention of QKI. Total QKI protein levels were unchanged (Figure 5D, E), confirming that the redistribution reflects nuclear translocation rather than altered QKI expression.

Alternative splicing analysis of LINC00941-depleted H1299 cells using rMATS [21] identified 3,233 statistically significant differential alternative splicing events (Figure 5F, G). These splicing changes are largely distinct from transcriptional changes identified by differential expression analysis, reflecting the mechanistically independent consequences of nuclear QKI activity. Intersection with a curated set of 75 established QKI target transcripts revealed that 62 of 77 were significantly enriched among the differentially spliced events, representing a highly significant overrepresentation relative to background (Figure 5H, I). Consistent with this, QKI target enrichment was confirmed by Fisher’s exact test (Figure 5H; p-value=0.001).

To validate these findings at the level of a canonical QKI target, we examined *NUMB* exon 12, a well-established readout of QKI nuclear activity [22]. RIP-qPCR demonstrated that knockdown of LINC00941 significantly enhanced QKI association with *NUMB* pre-mRNA (*>*2-fold enrichment; Figure 5J). RT-PCR and immunoblotting confirmed a significant shift in NUMB isoform usage, with a marked increase in the exon-12-excluded isoform in siLINC00941 cells (Figure 5K).

Co-depletion of QKI substantially rescued the anti-proliferative effect of siLINC00941 *in vitro* (Figure 5L), demonstrating that growth arrest caused by LINC00941 loss is partly driven by aberrant nuclear QKI splicing activity. In xenograft experiments, LINC00941 knockdown significantly suppressed tumour growth relative to control, and co-depletion of QKI substantially reversed this suppression (Figure 5M, N), providing *in vivo* evidence that LINC00941 operates principally through cytoplasmic sequestration of QKI [22].

### Clinical significance and therapeutic targeting of LINC00941 with antisense oligonucleotides

To assess clinical relevance, we interrogated three independent patient cohorts—TCGA, Michigan, and EAC—encompassing 649 patients in total. Univariate Cox regression analysis demonstrated that high LINC00941 expression was significantly associated with reduced overall survival in each cohort independently (TCGA: HR = 1.13, 95% CI 1.07–1.19, *p* = 9.34 × 10^−6^; Michigan: HR = 1.22, 95% CI 1.07–1.39, *p* = 3.38 × 10^−3^; EAC: HR = 1.02, 95% CI 1.00–1.03, *p* = 0.03; Figure 6B). Multivariate Cox regression incorporating patient stage as a covariate confirmed that LINC00941 remained an independent prognostic factor (TCGA: HR = 1.12, 95% CI 1.06–1.12, *p* = 3.13 × 10^−5^; Michigan: HR = 1.22, 95% CI 1.06–1.41, *p* = 5.92 × 10^−3^; EAC: HR = 1.01, 95% CI 1.00–1.03, *p* = 0.03; Figure 6B). Kaplan–Meier analysis further corroborated these findings (Figure 6C), establishing LINC00941 as an independent prognostic biomarker for poor outcome in LUAD.

**Figure 6.**
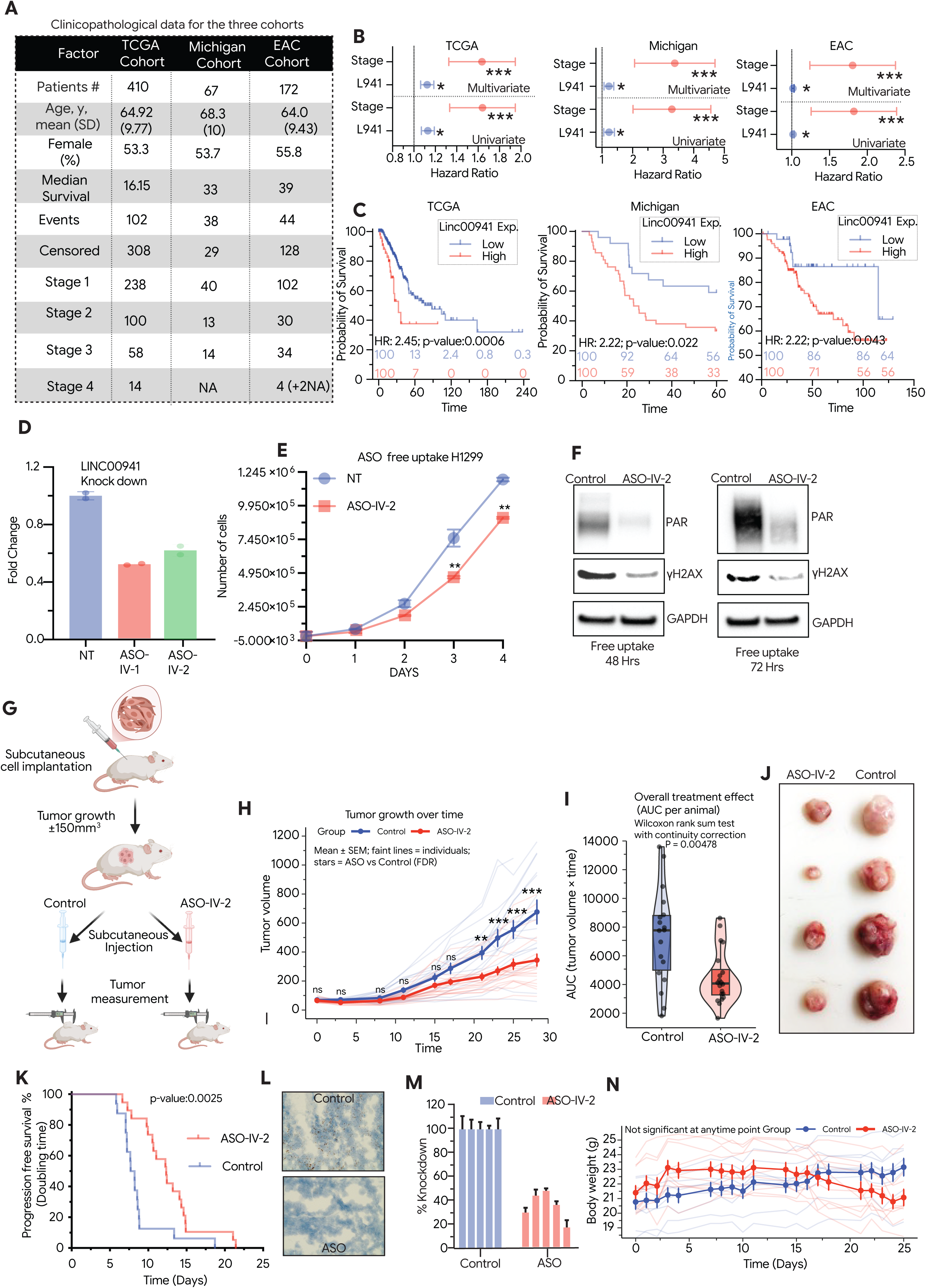
LINC00941 is a prognostic marker and a therapeutic vulnerability in lung adenocarcinoma. **(A)** Clinicopathological characteristics of patients included in the TCGA, MC, and East Asian lung adenocarcinoma cohorts. A total of 649 patients were analysed across all cohorts. **(B)** Univariate and multivariate Cox proportional hazards regression analyses performed in the TCGA, Michigan, and East Asian cohorts. LINC00941 expression emerged as a significant independent predictor of overall survival in all three datasets. **(C)** Kaplan–Meier survival analysis of patients stratified into high and low LINC00941 expression groups in the TCGA, Michigan, and East Asian cohorts. High LINC00941 expression was consistently associated with significantly poorer overall survival across all cohorts. **(D)** Quantitative RT-PCR analysis showing LINC00941 expression levels in cells treated with two independent antisense oligonucleotides (ASOs) targeting LINC00941 using a free-uptake method. ASO1 achieved higher knockdown of LINC00941. **(E)** Cell proliferation assay in H1299 cells following free-uptake treatment with control ASO or LINC00941-targeting ASO, demonstrating a significant reduction in cell proliferation upon LINC00941 depletion. **(F)** Analysis of DNA damage response markers following free-uptake ASO treatment. LINC00941 knockdown resulted in a significant decrease in PARylation levels and *γ*H2AX levels, consistent with results observed in siRNA- and shRNA-mediated knockdown experiments. **(G)** Schematic representation of the *in vivo* xenograft experimental design. H1299 cells were implanted subcutaneously into NOD/SCID mice, tumours were allowed to grow to approximately 150 mm^3^, and mice were subsequently treated with control or LINC00941-targeting ASOs. **(H)** Tumour growth curves showing tumour volume measurements over time in control- and ASO-treated groups. LINC00941 ASO treatment significantly suppressed tumour growth compared with control treatment. **(I)** Area-under-the-curve (AUC) analysis summarising the overall treatment effect, demonstrating a significant reduction in tumour burden over time in ASO-treated mice. **(J)** Representative images of tumours excised from control and ASO-treated mice at the experimental endpoint. **(K)** Progression-free survival analysis using tumour doubling time as the endpoint, showing significantly improved progression-free survival in ASO-treated tumours compared with controls. **(L)** ISH analysis of xenograft tumour sections showing reduced LINC00941 expression following ASO treatment. **(M)** Quantification of IHC staining demonstrating effective knockdown of LINC00941 in ASO-treated tumours. **(N)** Body weight measurements of mice throughout the treatment period, showing no significant differences between control and ASO-treated groups, indicating minimal systemic toxicity. Data are presented as mean ± SEM unless otherwise indicated. Statistical significance was determined using appropriate statistical tests as described in the Methods. ^∗^*P <* 0.05; ^∗∗^*P <* 0.01; ^∗∗∗^*P <* 0.001.

Given the consistent pro-tumorigenic role of LINC00941, we evaluated its potential as a therapeutic target. Two therapeutic-grade ASOs (ASO-IV-1 and ASO-IV-2; these are different ASOs than those used in figure 2C) targeting distinct regions of LINC00941 were designed and assessed for gymnotic uptake—free internalization without transfection reagents—in H1299 cells [23]. Both ASOs (ASO-IV-1 and ASO-IV-2) were efficiently internalised and achieved significant LINC00941 knockdown, with comparable efficacy (∼50% reduction; Figure 6D). ASO-mediated depletion significantly reduced cell proliferation (Figure 6E) and impaired DDR activity, as evidenced by decreased *γ*H2AX and PAR levels (Figure 6F). ASO-IV-2 demonstrated stronger *in vitro* performance and a superior systemic safety profile compared to ASO-IV-1, with markedly attenuated hepatic transaminase elevation and no organ weight abnormalities (Supplementary Figure 4). ASO-IV-2 was therefore selected for all subsequent in vivo studies.

Once tumours reached a mean volume of ∼150 mm^3^, mice were randomised (control *n* = 18; ASO-IV-2 *n* = 20) and dosed twice weekly for four weeks. ASO-IV-2 treatment resulted in significant suppression of tumour growth, with mean tumour volumes at endpoint of 750 ± 300 mm^3^ (control) versus 359 ± 193 mm^3^ (ASO-IV-2-treated; *p <* 0.001; Figure 6H, I, J). Kaplan–Meier analysis of tumour volume doubling time (analogous to progression-free survival) revealed significantly prolonged progression-free survival in ASO-treated mice (median: 7.69 days control vs. 12.35 days ASO-IV-2; log-rank *p* = 0.002; Figure 6K). Post-treatment molecular analysis confirmed successful *in vivo* target engagement by RNA-ISH and RT-qPCR (Figure 6L, M). ASO-IV-2 treatment was well tolerated with no significant effect on body weight (Figure 6M). These findings provide robust preclinical proof-of-concept that gymnotic ASO-mediated targeting of LINC00941 effectively suppresses tumour growth *in vivo* [24, 25].

## Discussion

KRAS mutations are the most prevalent oncogenic driver in LUAD, occurring in approximately 30% of cases, yet the transcriptional programmes downstream of KRAS-MAPK signalling that sustain cancer cell fitness remain incompletely characterised. The AP-1 transcription factor FOSL1 (FRA-1) is a well-established effector of RAS-ERK signalling, and its overexpression in multiple tumour types has been linked to proliferation, invasion, and therapy resistance [26]. Despite extensive characterisation of FOSL1 protein-coding targets, the lncRNA transcriptional programme controlled by FOSL1 downstream of KRAS-MAPK in lung cancer has not previously been described. Here we demonstrate that LINC00941 is a direct transcriptional target of FOSL1, establishing what is, to our knowledge, the first FOSL1-regulated lncRNA in the KRAS-MAPK pathway. LINC00941 expression correlated with FOSL1 levels across LUAD datasets regardless of KRAS mutation status, though KRAS-mutant cells exhibited consistently higher expression, suggesting that while FOSL1-dependent transcription is the primary driver, additional KRAS-independent inputs may contribute to LINC00941 regulation. It is important to note that while the discovery experiment was performed in A549 cells harbouring KRAS^G12S^, a key mechanistic cell model throughout this study—H1299—carries an NRAS^Q61K^ mutation. Both mutations constitutively activate the RAF–MEK–ERK cascade and induce equivalent FOSL1 upregulation, reflecting the pan-RAS nature of this transcription factor. The consistent regulation and function of LINC00941 observed across both cell lines indicates that the LINC00941 regulatory axis is a general consequence of oncogenic RAS–MAPK–FOSL1 hyperactivation rather than a *KRAS* allele-specific phenomenon. This broader RAS pathway relevance is a strength of the present work: whereas KRAS is the dominant driver in LUAD (25–30% of cases) and provides the primary clinical hook, the shared regulation of LINC00941 by both KRAS and NRAS oncoproteins through a common FOSL1-dependent mechanism suggests that the findings may extend to RAS-driven malignancies beyond lung adenocarcinoma.

The primary conceptual contribution of this work is the demonstration that LINC00941 functions through two mechanistically distinct, compartment-specific axes that collectively provide replication stress tolerance and splicing fidelity to LUAD cells. This dual-compartment regulatory model advances an emerging paradigm in lncRNA biology—that subcellular localisation is not merely a static property but a deterministic feature that divides a single lncRNA into functionally non-overlapping molecular entities operating in parallel [10]. In the nucleus, LINC00941 physically associates with SAM68 (KHDRBS1) through its 700–1300 nucleotide region and shields it from proteasomal degradation. SAM68 is an essential co-activator of PARP1-mediated poly(ADP-ribosyl)ation—a critical early event in the DNA damage response—and its genetic deletion results in dramatically impaired PAR synthesis and hypersensitivity to genotoxic stress [20]. Our findings extend this foundational work by demonstrating that SAM68 protein stability itself is regulated by a lncRNA in a cancer-specific context. The observation that LINC00941 depletion phenocopies SAM68 loss—with abrogated pCHK1 signalling, reduced PAR synthesis, impaired *γ*H2AX and 53BP1 foci formation, and increased comet tail moment—and that SAM68 overexpression rescues these defects, places LINC00941 upstream of an established DDR axis as a novel stability regulator. Importantly, the accelerated proteasomal turnover of SAM68 in LINC00941-depleted cells, rescued by MG132, suggests a model in which LINC00941 binding sterically occludes proteosomal targeting of SAM68. The identification of RNF123 as the candidate E3 ubiquitin ligase mediating SAM68 degradation is supported by its consistent negative correlation with SAM68 protein levels across two independent datasets — the Cancer Cell Line Encyclopedia and the CPTAC LUAD proteome — and by the functional observation that RNF123 co-depletion rescues SAM68 protein levels in LINC00941-knockdown cells. We acknowledge that direct biochemical demonstration of RNF123-mediated SAM68 ubiquitination — including physical interaction between RNF123 and SAM68 and RNF123-dependent polyubiquitin chain formation — remains to be established, and that the current evidence is correlational and functional rather than biochemically complete. Nevertheless, the convergence of cross-dataset proteogenomic correlation with functional rescue provides sufficient grounds to nominate RNF123 as the principal E3 ligase governing LINC00941-dependent SAM68 stability, a mechanistic relationship that warrants direct biochemical interrogation in future work. A recent study in breast cancer stem cells further supports the therapeutic relevance of the SAM68–PARP1 axis, demonstrating synthetic lethality between SAM68 inhibition and RAD51 suppression [27], raising the possibility that LINC00941-high tumours may exhibit differential sensitivity to PARP inhibitors.

In the cytoplasm, LINC00941 acts through an orthogonal mechanism by sequestering QKI, preventing its nuclear translocation and consequent aberrant splicing of target pre-mRNAs. QKI is an established splicing regulator and tumour suppressor in NSCLC, where reduced nuclear QKI expression correlates with worse disease-free survival [22]. Paradoxically, our data show that in LINC00941-expressing LUAD cells, cytoplasmic retention of QKI is pro-tumorigenic—its nuclear translocation upon LINC00941 loss drives widespread splicing dysregulation affecting 24 of 75 known QKI target genes. The *NUMB* exon 12 splicing shift we validate is particularly instructive: QKI-dependent NUMB isoform switching attenuates Notch signalling, which in this context impairs rather than reinforces cancer cell proliferative signalling. The *in vitro* and *in vivo* rescue of the anti-proliferative phenotype by QKI co-depletion provides epistatic evidence that unrestrained nuclear QKI activity is the functional mediator of growth arrest upon LINC00941 loss—a finding that reframes QKI’s role as context- and localisation-dependent, consistent with the emerging understanding that lncRNA-mediated subcellular sequestration of RBPs can fundamentally alter their functional output [10].

The dual-compartment architecture of LINC00941 raises the question of how a lncRNA with approximately equal nuclear and cytoplasmic residency executes mechanistically distinct functions in each compartment without cross-interference. We propose that this is an intrinsic consequence of the compartment-restricted availability of its binding partners: SAM68 is a predominantly nuclear protein and its interaction with LINC00941 is therefore confined to the nuclear fraction, while QKI—though capable of nucleocytoplasmic shuttling—is retained in the cytoplasm under basal conditions precisely through its engagement with cytoplasmic LINC00941. In this model, the nuclear and cytoplasmic pools of LINC00941 function as molecularly distinct entities whose activities are determined not by the lncRNA sequence *per se* but by the local protein environment of each compartment. This framework is consistent with the emerging understanding that subcellular localisation acts as a deterministic switch that partitions lncRNA function across compartments [10], and is directly supported by our compartment-resolved RIP data showing preferential SAM68 enrichment in the nuclear fraction and preferential QKI enrichment in the cytoplasmic fraction. While the two mechanisms operate in parallel rather than in series—as evidenced by the independent epistatic rescue experiments for each arm—they converge on the same tumour-promoting outcome: maintenance of replication stress tolerance in KRAS-driven LUAD cells. Whether dynamic redistribution of LINC00941 between compartments in response to oncogenic stress signals further fine-tunes the balance between these two activities remains an important question for future investigation.

Taken together, both nuclear and cytoplasmic mechanisms converge on the same tumour-promoting outcome: maintenance of DDR competency and splicing fidelity under replicative stress. Cancer cells harbouring oncogenic KRAS face elevated replication stress and therefore possess a heightened dependency on DDR pathways. Our data suggest that LINC00941 is a molecular solution co-opted by KRAS-driven cancer cells to sustain this dependency: by simultaneously stabilising SAM68 to maintain PARP1 activity and sequestering QKI to preserve splicing of DNA repair transcripts including *RAD51*, *BRCA2*, and *FANCD2*, LINC00941 provides a dual layer of protection against replication stress-induced cytotoxicity. This framework resolves the apparent paradox of an oncogenic lncRNA that, when depleted, induces both increased DNA damage and impaired DDR signalling—the two phenotypes are causally linked through the SAM68–PARP1 axis.

The clinical and therapeutic implications of this work are compelling. Multi-cohort Cox regression across TCGA, Michigan, and EAC LUAD datasets demonstrates that LINC00941 is an independent prognostic factor for poor survival after adjustment for stage. More significantly, gymnotic delivery of ASO-IV-2 achieved potent *in vivo* target knockdown and significantly suppressed xenograft growth at *p* = 0.002, with no measurable systemic toxicity. Gymnotic ASO uptake is a clinically relevant delivery modality [23], and the precedent for ASO-mediated targeting of oncogenic lncRNAs *in vivo* is growing [24, 25], providing a conceptual framework for the translational approach undertaken here.

Several limitations of the present study merit acknowledgement. The transcriptomic analysis was conducted with two independent siRNAs, each represented by a single sequencing lane, which limits formal statistical power for differential expression analysis; the parallel microarray dataset provides important cross-platform validation, though the RNA-seq findings should be interpreted as hypothesis-generating at the individual gene level. Mechanistic studies were conducted primarily in H1299 and A549 cell lines, and validation in patient-derived models or additional NSCLC lines would strengthen generalisability. The precise ubiquitination site and further validation of RNF123 mediated SAM68 degradation in the absence of LINC00941 remain to be fully determined, as does the structural basis of the QRE-mediated LINC00941–QKI interaction. Future work exploring whether LINC00941 expression modulates sensitivity to PARP inhibitors or genotoxic chemotherapy in KRAS-mutant LUAD would be of direct therapeutic relevance, as would investigation of whether the LINC00941–SAM68 axis constitutes a synthetic lethal vulnerability in BRCA-deficient contexts [27].

In conclusion, this study establishes LINC00941 as a novel oncogenic lncRNA in LUAD that operates through a dual-compartment mechanism—nuclear stabilisation of SAM68 to sustain DDR competency, and cytoplasmic sequestration of QKI to preserve splicing fidelity of DNA repair transcripts—both converging to provide replication stress tolerance in KRAS-driven lung cancer. The identification of LINC00941 as the first FOSL1-regulated lncRNA, combined with its independent prognostic value and preclinical ASO targetability, positions it as both a mechanistically informative node in the KRAS-MAPK transcriptional network and a tractable therapeutic target in LUAD.

## Materials and Methods

### Cell lines and culture conditions

Human non-small cell lung cancer (NSCLC) cell lines H1299, H358, H23, and H1437 were obtained from the American Type Culture Collection (ATCC, USA); A549 was obtained from the National Centre for Cell Science (NCCS; Pune, India). All cell lines were maintained in recommended media supplemented with 10% foetal bovine serum (FBS) and 1% penicillin–streptomycin at 37 ^◦^C in a humidified atmosphere containing 5% CO_2_. Cell lines were genotyped to confirm identity by short tandem repeat (STR) profiling performed at NCCS, Pune. Cells line culture were routinely tested for *Mycoplasma* contamination by PCR using the EZdetect™ PCR kit (HiMedia; Cat. No. CCK022) and only mycoplasma-free cultures were used for experiments.

### siRNA transfection

Small interfering RNA (siRNA)-mediated knockdown was performed using Lipofectamine RNAiMAX transfection reagent (Thermo Fisher Scientific) according to the manufacturer’s protocol. Cells were seeded in antibiotic-free medium and transfected with siRNA at a final concentration of 30 nM. Knockdown efficiency was assessed by quantitative RT-PCR and/or western blotting 48– 72 hours post-transfection as appropriate. All siRNA sequences are listed in Supplementary Table ST1.

### Lentiviral production and generation of stable cell lines

Lentiviral particles were produced using the Lenti-X packaging system (Takara Bio, Japan) in LentiX packaging cells. LentiX cells were co-transfected with the shRNA expression vector and packaging plasmids using X-fect transfection reagent (Takara Bio, Japan Bio, Japan) according to the manufacturer’s instructions. Virus-containing supernatants were harvested at 48 and 72 hours post-transfection, filtered through 0.45 *µ*m membranes, and concentrated using LentiX concentrator (Takara Bio, Japan Bio, Japan). Target cells were transduced in the presence of 8 *µ*g/ml polybrene (Sigma-Aldrich) and stable integrants were selected with puromycin (2–4 *µ*g/ml) for 2–4 days. Knockdown efficiency was verified by RT-PCR and western blot prior to downstream experiments. shRNA sequences are listed in Supplementary Table ST1.

### Antisense oligonucleotide (ASO) transfection

Two independent antisense oligonucleotides (ASOs, IDT, USA) targeting LINC00941 and a matched non-targeting control were used for orthogonal validation of siRNA phenotypes. ASOs were transfected using Lipofectamine RNAiMAX at a final concentration of 30 nM following the same protocol described above. Knockdown efficiency was confirmed by RT-PCR 48–72 hours post-transfection. ASO sequences and chemical modifications are provided in Supplementary Table ST1.

### RNA extraction and quantitative RT-PCR

Total RNA was extracted using the RNeasy Mini Kit (Qiagen) according to the manufacturer’s instructions. RNA quantity and integrity were assessed by NanoDrop spectrophotometry and agarose gel electrophoresis, respectively. Complementary DNA (cDNA) was synthesised from 500 ng–1 *µ*g total RNA using the PrimeScript™ 1st-strand cDNA Synthesis Kit (Takara Bio, Japan). Quantitative PCR was performed on a Bio-Rad qPCR system using SYBR Green Master Mix (Thermo Scientific). Relative expression was calculated by the 2^−ΔΔCt^ method and normalised to either *GAPDH* and/or *TPT1* as internal reference genes. All primer sequences are listed in Supplementary Table ST2.

### RNA sequencing and bioinformatic analysis

Two independent RNA-seq experiments were performed. First, to identify genes regulated by KRAS–MAPK signalling, A549 cells were treated with the MEK1/2 inhibitor Cobimetinib (1 µM, 48 h) or DMSO vehicle control (*n* = 2 biological replicates per condition). Second, to characterise the global transcriptional consequences of LINC00941 loss, H1299 cells were transfected with siLINC00941 (*n* = 2 biological replicates) or non-targeting siRNA control (siNT; *n* = 4 biological replicates) and harvested 72 hours post-transfection. For both experiments, total RNA was extracted using RNAeasy kit (Qiagen, USA) and RNA quality was assessed using a Bioanalyser (Agilent) with a minimum RNA Integrity Number (RIN) ≥ 8.0 required for library preparation. Strandspecific RNA-seq libraries were prepared using poly-A enrichment. Libraries were sequenced on the Illumina NovaSeq platform (SNPcode, Mumbai, India) with paired-end 150 bp reads to a minimum depth of 30 million reads per sample.

Raw reads were quality-assessed with FastQC [28] and adapter-trimmed using Trimmomatic [29]. Reads were aligned to the human reference genome (GRCh38/hg38) using STAR (v2.5.2b), and GENCODE gene annotations [30]. Gene-level read counts were quantified with featureCounts [31] against the GENCODE vGRCh38.p14 annotation. Differential expression analysis was performed in R (v4.x) using DESeq2 [32], with default shrinkage estimators applied. Genes with a base mean FPKM *>* 0.5 in at least one group were retained. Differentially expressed genes (DEGs) were defined by adjusted *p <* 0.05 (Benjamini–Hochberg correction) and | log_2_ FC| *>* 1. Pathway enrichment analysis was performed by manual gene-set classification against KEGG pathway annotations [33, 34]. Alternative splicing analysis was performed using rMATS [21]. QKI target gene enrichment was assessed by Fisher’s exact test using a curated list of 75 established QKI target genes [22]. All the Sequencing data is submitted as Supplementary Table ST3.

### Western blotting

Cells were lysed in RIPA buffer (50 mM Tris-HCl pH 8.0, 150 mM NaCl, 1% NP-40, 0.5% sodium deoxycholate, 0.1% SDS) supplemented with protease and phosphatase inhibitor cocktails (Sigma-Aldrich). Protein concentrations were determined by BCA assay (Pierce). Equal quantities of protein (20–40 *µ*g) were resolved by SDS-PAGE on 8–12% polyacrylamide gels and transferred to PVDF membranes (Millipore). Membranes were blocked in 5% non-fat dried milk or 5% BSA in TBST for 1 hour at room temperature, then incubated with primary antibodies overnight at 4 ^◦^C. After washing, membranes were incubated with HRP-conjugated secondary antibodies for 1 hour at room temperature. Bands were visualised by enhanced chemiluminescence (ECL) on a Bio-Rad imaging system. *β*-Actin or GAPDH was used as a loading control. All antibodies are listed in **Supplementary Table ST3**.

### Immunofluorescence and confocal microscopy

Cells grown on glass coverslips were fixed in 4% paraformaldehyde (PFA) for 15 minutes at room temperature and permeabilised with 0.3% Triton X-100 in PBS for 10 minutes. After blocking in 5% normal goat serum (NGS) in PBS for 1 hour, coverslips were incubated with primary antibodies overnight at 4 ^◦^C. Cells were washed and incubated with Alexa Fluor-conjugated secondary antibodies (1:500; Thermo Fisher Scientific) for 1 hour at room temperature in the dark. Nuclei were counterstained with DAPI (0.1 *µ*g/ml). Images were acquired on a Zeiss confocal laser scanning microscope. Fluorescent foci (*γ*H2AX, 53BP1) were quantified in a minimum of 50 cells per condition across three independent experiments using ImageJ/Fiji [35].

### Chromatin immunoprecipitation (ChIP-qPCR)

Cells (∼10×10^6^) were cross-linked with 1% formaldehyde for 10 minutes at room temperature, quenched with 125 mM glycine, and harvested. Chromatin was isolated and sonicated to an average fragment size of 200–500 bp using a Diagenode Bioruptor Plus. Immunoprecipitation was performed overnight at 4 ^◦^C with anti-FOSL1 antibody or normal rabbit IgG as a negative control, using protein A/G magnetic beads (Thermo Fisher Scientific). After serial washes, cross-links were reversed at 65 ^◦^C overnight and DNA was purified using a PCR purification kit (Qiagen). Enrichment of LINC00941 promoter regions was quantified by qPCR and expressed as a enrichment over IgG. Primer sequences are listed in Supplementary Table ST5.

### RNA immunoprecipitation (RIP)

RIP was performed using the RIP-Assay Kit (MBL, Japan) supplemented with RNase inhibitor and protease inhibitors. Lysates were pre-cleared with protein A/G magnetic beads, then incubated with anti-SAM68, anti-QKI, or control IgG overnight at 4 ^◦^C. Co-immunoprecipitated RNA was isolated using TRIzol LS, reverse transcribed, and LINC00941 enrichment was quantified by qPCR and expressed as fold enrichment over IgG control. Antibody details are provided in Supplementary Table ST3.

### RNA pulldown assay

Full-length LINC00941 and control antisense RNA were transcribed *in vitro* from linearised plasmid templates using T7 RNA polymerase (Thermo Fisher Scientific). RNA was biotinylated using the Pierce™ RNA 3’ End Biotinylation Kit. Biotinylated RNA (5 *µ*g) was folded in RNA structure buffer and incubated with pre-cleared cell lysates for 2 hours at room temperature. Streptavidin-conjugated magnetic beads (Thermo Fisher Scientific) were added for a further 30 minutes. Bound proteins were eluted, resolved by SDS-PAGE, and analysed by western blotting for SAM68 and QKI.

### Subcellular fractionation

Nuclear and cytoplasmic fractions were separated using the NE-PER Nuclear and Cytoplasmic Extraction Reagents (Thermo Fisher Scientific) according to the manufacturer’s protocol. Fraction purity was verified by western blotting using GAPDH (cytoplasmic marker). RNA from each fraction was extracted and used for qRT-PCR to determine LINC00941 subcellular distribution.

### Alkaline comet assay

DNA strand break accumulation was quantified by alkaline single-cell gel electrophoresis (comet assay) using the CometAssay Kit (R&D Systems). Cells were harvested 72 hours post-transfection/infection and embedded in 0.5% low-melting-point agarose at ∼10,000 cells/ml. Slides were lysed in alkaline lysis buffer for 1 hour at 4 ^◦^C in the dark, then subjected to alkaline electrophoresis (300 mM NaOH, 1 mM EDTA, pH *>* 13) at 25 V and 300 mA for 30 minutes. Slides were neutralised, fixed in ice-cold 70% ethanol, and stained. Comet tail length was quantified from a minimum of 50 cells per condition using ImageJ. Comet images were processed/modified using CometScore. A minimum of three independent biological replicates were performed.

### Cell proliferation and colony formation assays

Cell proliferation was assessed by direct cell counting using a LUNA 2 automated cell counter at 24, 48, 72, and 96 hours post-transfection, and independently monitored using the Incucyte live-cell imaging system. For colony formation, 500–1000 cells per well were seeded in six-well plates and cultured for 10–14 days. Colonies were fixed with methanol and stained with 0.5% crystal violet in 25% methanol, then quantified using ImageJ. All assays were performed in triplicate across at least three independent experiments.

### Senescence-associated *β*-galactosidase ((SA-*β*-Gal) staining

Cellular senescence was assessed using the Senescence *β*-Galactosidase Staining Kit (Cell Signaling Technology, #9860) according to the manufacturer’s instructions. SA-*β*-Gal-positive cells (blue staining at pH 6.0) were quantified by bright-field microscopy from a minimum of 200 cells per condition across three independent experiments.

### Cell cycle analysis

Cells were harvested 72 hours post-transfection, washed with PBS, and fixed in ice-cold 70% ethanol overnight at −20 ^◦^C. Fixed cells were resuspended in PI staining solution (50 *µ*g/ml propidium iodide, 100 *µ*g/ml RNase A, 0.1% Triton X-100 in PBS) and incubated for 30 minutes at 37 ^◦^C in the dark. DNA content was measured by flow cytometry with a minimum of 10,000 events recorded per sample. Cell cycle phase distributions were calculated using ModFit software.

### In vivo xenograft tumour model

All animal experiments were performed in accordance with Institutional Animal Ethics Committee (IAEC) guidelines and were approved under the relevant protocols. Sixto eight-week-old NOD-SCID mice were used for subcutaneous xenograft experiments. H1299 cells stably expressing shRNA against LINC00941, QKI, both targets, or non-targeting shRNA control (shNT) were resuspended in a 1:1 mixture of PBS and Matrigel (Corning). One million (1 × 10^6^) cells per mouse were injected subcutaneously into the right flank. Tumour dimensions were monitored by caliper measurement every two to three days and volumes calculated as: *V* = (length × width^2^)*/*2. Animals were sacrificed after four weeks or when tumours reached the ethical endpoint of 1,500 mm^3^. Tumours were excised, weighed, and processed for histological and molecular analyses.

### In vivo ASO treatment

NOD-SCID mice bearing established subcutaneous H1299 xenografts were randomised into vehicle control (*n* = 18) and ASO-IV-2 treatment (*n* = 20) groups once tumours reached a mean volume of ∼150 mm^3^. ASO-1 was formulated in sterile PBS at 12.5 mg/ml and administered subcutaneously at 50 mg/kg twice weekly for four weeks, with injection volume adjusted for individual body weight. Dosing volumes for the expected weight range (15–29 g) are provided in Supplementary Table ST6. Tumour volumes were measured as described above. At study termination, tumours were excised and processed for RNA and protein analyses. Body weights were recorded throughout as a measure of systemic tolerability. Gymnotic ASO delivery in the absence of transfection reagents was assessed as described previously [23].

### Statistical analysis

All statistical analyses were performed in R (version 4.x; R Core Team) or GraphPad Prism version 10. Unless otherwise stated, data are presented as mean ± standard deviation (s.d.) or mean ± standard error of the mean (s.e.m.) from a minimum of three independent biological experiments. Comparisons between two groups used two-tailed Student’s *t*-test (normally distributed data) or Mann–Whitney *U* -test (non-parametric data). Multiple group comparisons used one-way ANOVA with Tukey’s post-hoc test. Fisher’s exact test (one-tailed) was used for gene-set enrichment analysis of QKI target genes within DEG lists. For *in vivo* tumour growth curves, statistical significance was assessed using two-way ANOVA with Bonferroni correction. Differential expression analysis used the Benjamini–Hochberg procedure for multiple testing correction. *p <* 0.05 was considered statistically significant throughout.

## Acknowledgements

Sudhanshu Shukla acknowledges funding from the Indian Council for Medical Research, Government of India (Grant No. 2021-9513/CMB/ADHOC-BMS and EMDR/SG/13/2023-0244), the Department of Biotechnology, Government of India (Grant ID: BT/PR51308/MED/30/2511/2023), and the Anusandhan National Research Foundation, Government of India (Grant ID: CRG/2023/000837). Claude and Grammarly were used for paraphrasing and grammar correction. A.M.C. is a Howard Hughes Medical Institute Investigator, A. Alfred Taubman Scholar and American Cancer Society Professor.

## Author Contributions

**DA:** Methodology, Formal analysis, Investigation, Writing – Review and Editing.

**JCYT:** Animal Experiments.

**AS (Anu Sharma):** Investigation.

**VB:** Investigation.

**AS (Aaditya Singh):** Investigation.

**MA:** Investigation.

**SP:** Investigation.

**XC:** Investigation.

**RM:** Investigation.

**SMD:** Investigation.

**BKC:** Writing – Review and Editing, Supervision.

**AMC:** Conceptualization, Writing – Review and Editing, Funding Acquisition, Supervision.

**SS:** Conceptualization, Methodology, Formal Analysis, Writing – Review and Editing, Funding Acquisition, Supervision.

---

**Supplementary Figure 1.**
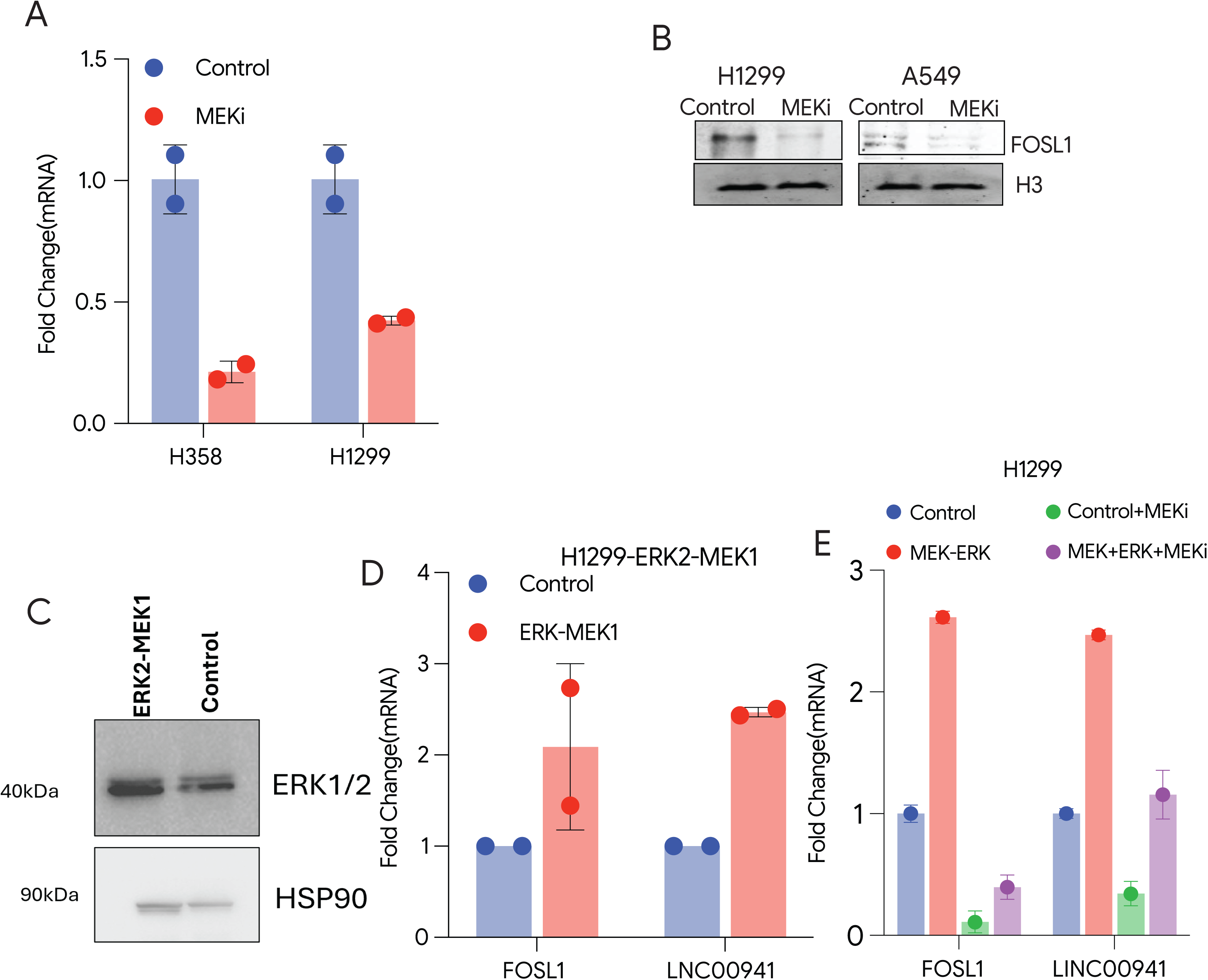
(A) q-RTPCR data of H1299 and H358 cells treated with MEKi. PCR was done for LINC00941. (B) Immunoblot analysis showing the effect of MEK inhibition on LINC00941 protein expression. Cells were treated with a MEK inhibitor, and FOSL1 levels were assessed by Western blotting. (C) Western blot data showning increased expression of ERK1/2 after overexpression in H1299 cells. (D) q-RTPCR data of H1299 treated with MEKi. PCR was done for FOSL1 and LINC00941. (E) Control or MEK-ERK fusion gene was overexpressed in H1299 cells and were treated with DMSO or MEK inhibitor, then qRT-PCR was done for FOSL1 and LINC00941-RNA.

**Supplementary Figure 2:**
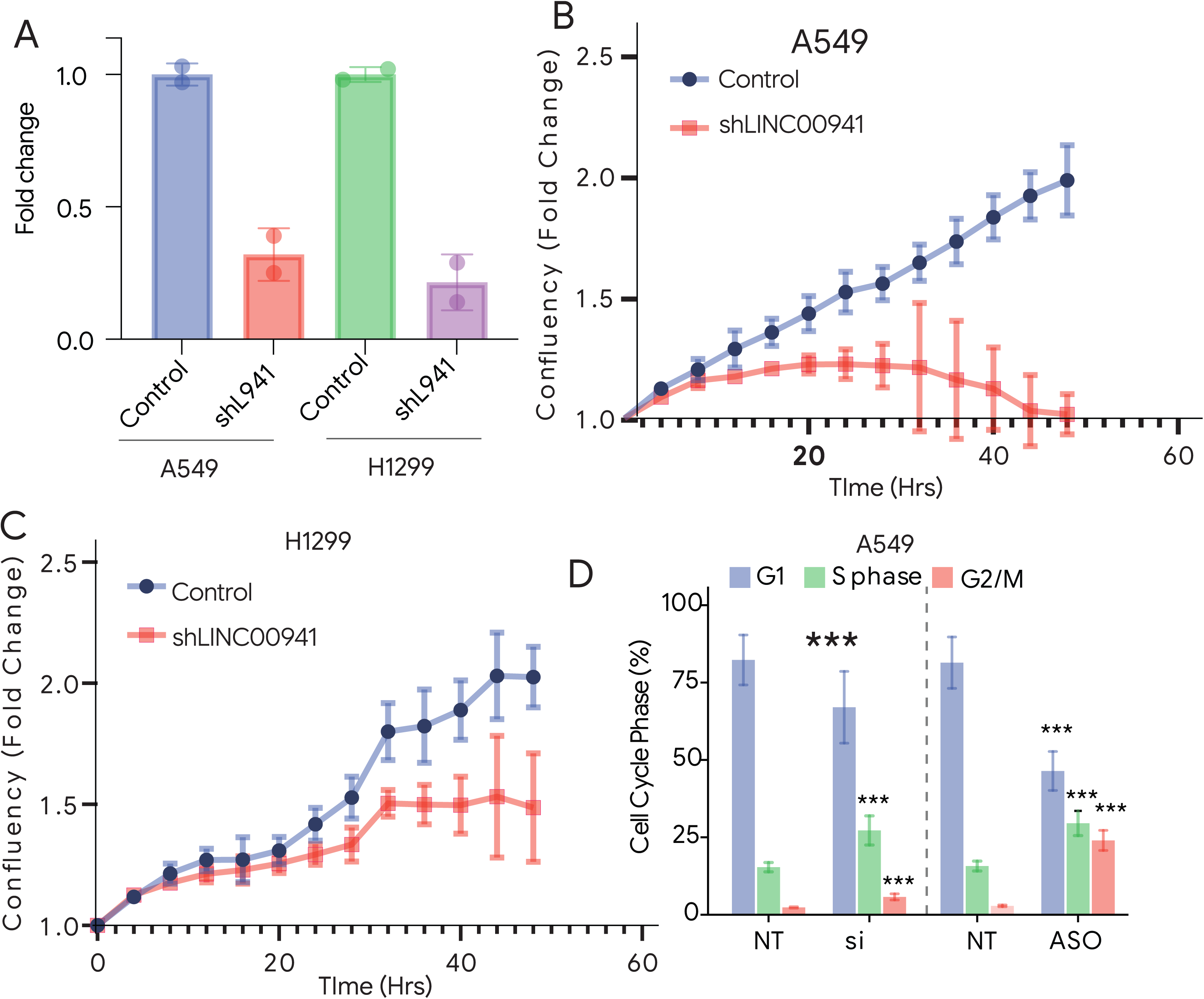
(A) q-RTPCR data of control and shLINC00941 in A549 and H1299 cells (B) Cell proliferation assay performed following LINC00941 knockdown using shRNAs, showing a significant decrease in proliferation in LINC00941 depleted H1299 cells compared with controls. (C) Cell proliferation assay performed following LINC00941 knockdown using shRNAs, showing a significant decrease in proliferation in LINC00941 depleted A549 cells compared with controls. (D) Cell cycle analysis of cells transfected with control or LINC00941 targeting siRNA. Flow cytometric profiling revealed enrichment of cells in S phase and G2/M phase following LINC00941 knockdown.

**Supplementary Figure 3.**
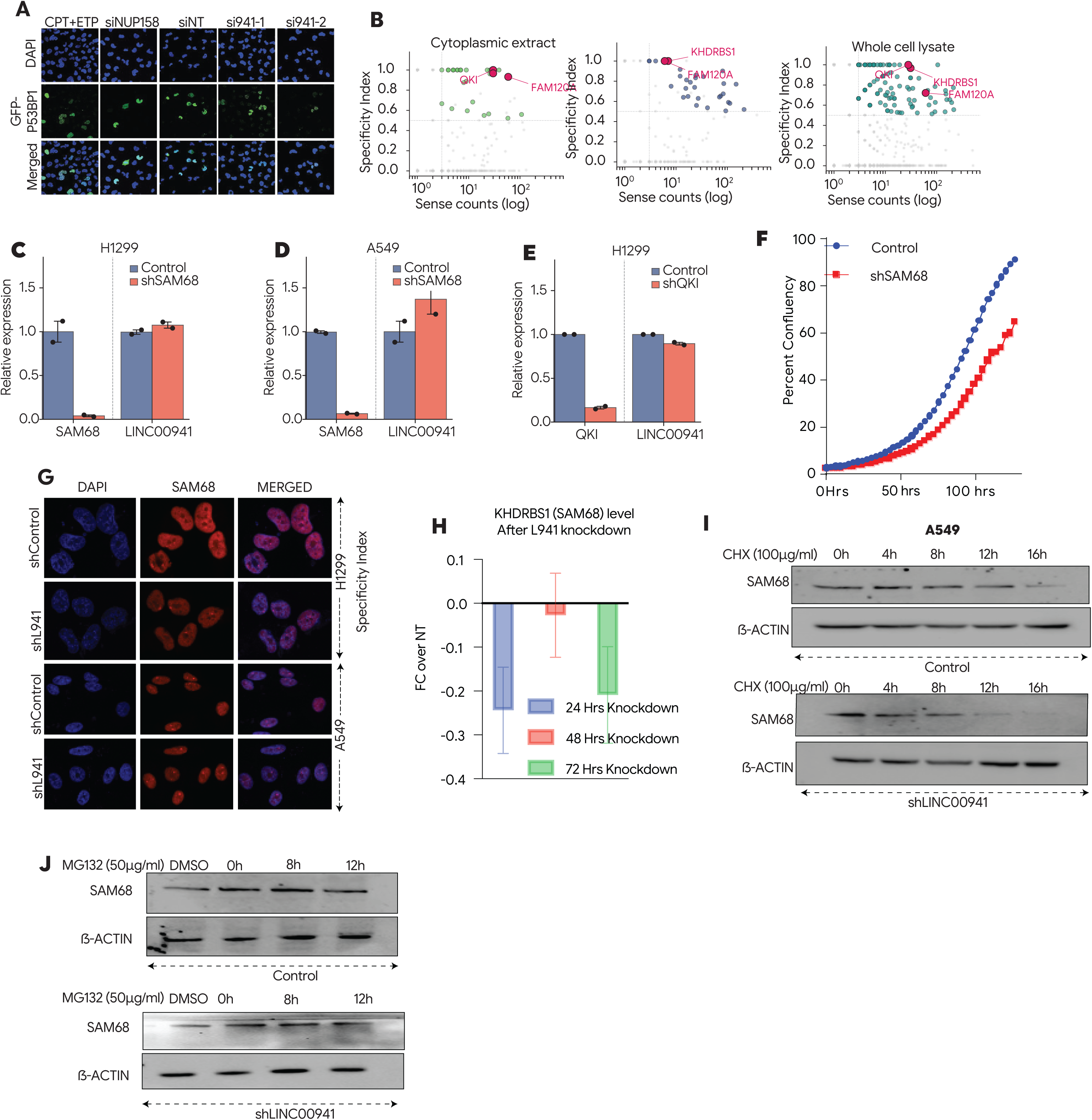
(A) Immunofluorescence analysis of p53BP1 foci in control and LINC00941 depleted cells. Representative images and quantification demonstrate a significant reduction in p53BP1 foci upon LINC00941 knockdown. (B) Scatter plots showing the enrichment of proteins identified in LINC00941 RNA pulldown mass spectrometry across cytoplasmic, nuclear, and whole cell lysate fractions. (C-E) RT-qPCR analysis of SAM68 and LINC00941 expression in (C) H1299 and (D) A549 cells stably expressing shSAM68 or a control shRNA. (E) RT-qPCR analysis of QKI and LINC00941 expression in H1299 cells expressing shQKI or a control shRNA. All values are normalised to control . (F) Cell proliferation assay performed following SAM68 knockdown, showing a significant decrease in proliferation in SAM68 depleted cells compared with controls. (G) IF analysis of SAM68 protein levels in control and LINC00941 knockdown cells, showing decresed level of total SAM68 expression upon LINC00941 depletion. (H) RNA level of of SAM68 gene after LINC00941 knockdown. (I) Cycloheximide chase assay assessing SAM68 protein stability in control and shLINC00941 A549 cells. Cells were harvested at the indicated time points following cycloheximide treatment and SAM68 protein levels were analysed by immunoblotting. (J) Proteasome inhibition assay showing stabilisation of SAM68 protein in control and shLINC00941 cells treated with MG132, indicating proteasome-mediated degradation of SAM68 upon LINC00941 depletion.

**Supplementary Figure 4:.**
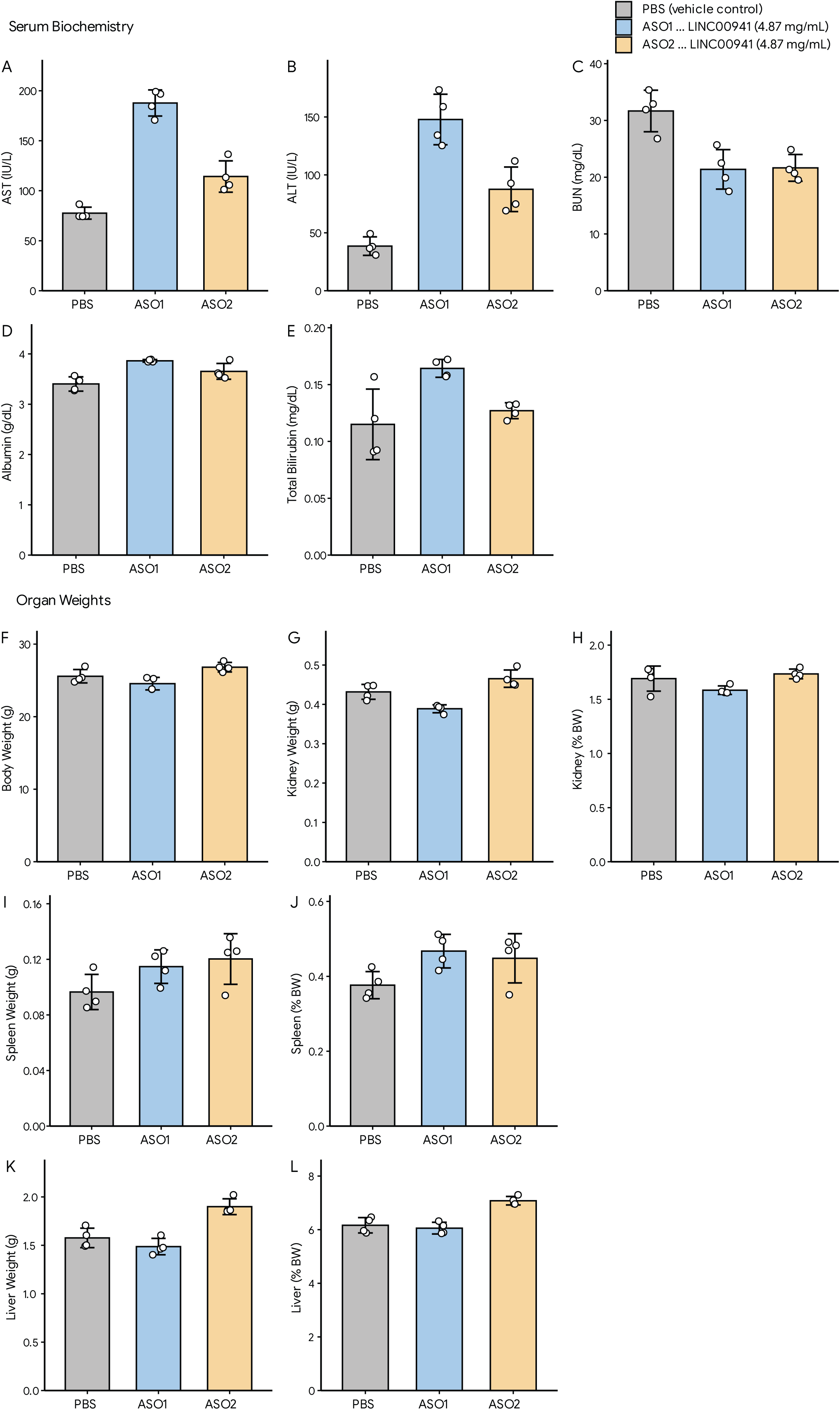
Systemic tolerability assessment of LINC00941-targeting antisense oligonucleotides ASO1 and ASO2 in vivo. Animals were administered PBS (vehicle control), ASO1, or ASO2 (both targeting LINC00941; 4.87 mg/mL, 100 mg/kg/week) and sacrificed at study endpoint for biochemical and morphometric analysis. (A) Serum aspartate aminotransferase (AST) levels. (B) Serum alanine aminotransferase (ALT) levels. (C) Blood urea nitrogen (BUN). (D) Serum albumin. (E) Total serum bilirubin. (F) Terminal body weights. (G, H) Kidney weight expressed as absolute mass (g) and relative to body weight (% BW), respectively. (I, J) Spleen weight expressed as absolute mass (g) and relative to body weight (% BW), respectively. (K, L) Liver weight expressed as absolute mass (g) and relative to body weight (% BW), respectively. Bars represent group mean ± SD; individual animal data points are overlaid. n = 4 per group. PBS, grey; ASO1, blue; ASO2, amber.

**Supplementary Table S1.**
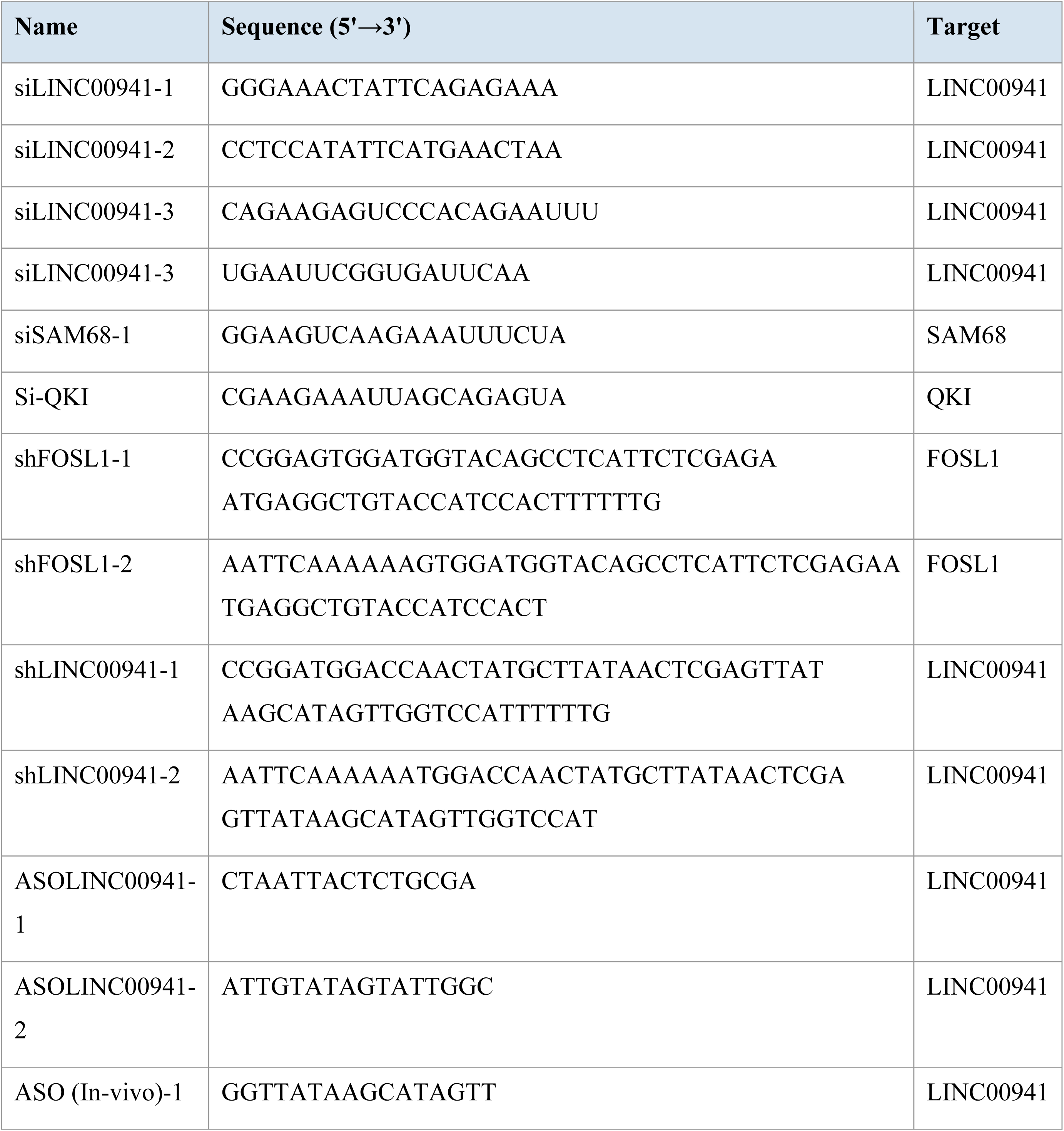
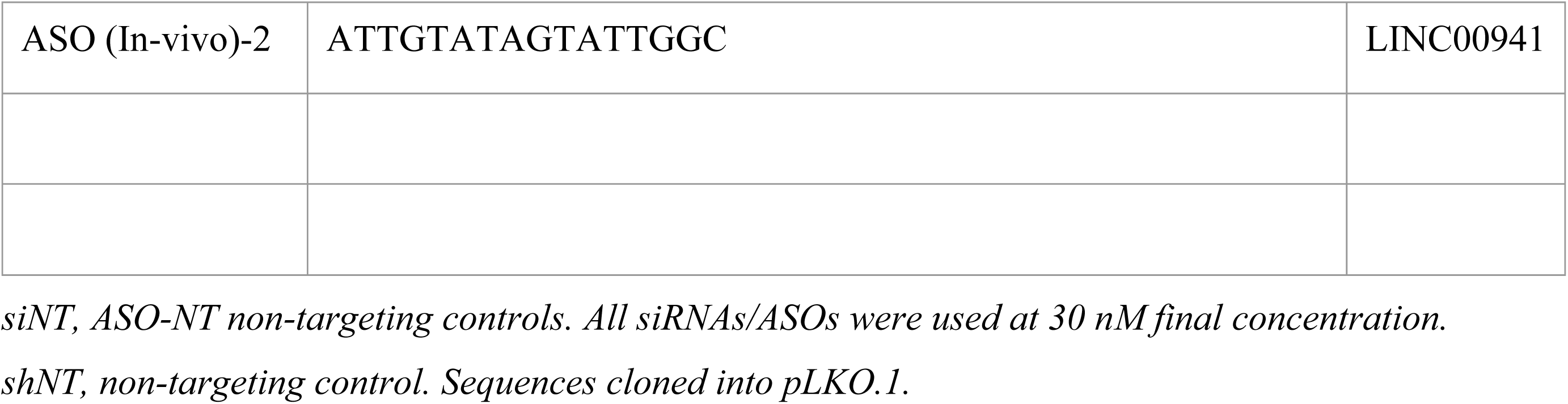
siRNA, ASO and shRNA sequences used in this study.

**Supplementary Table S2.**
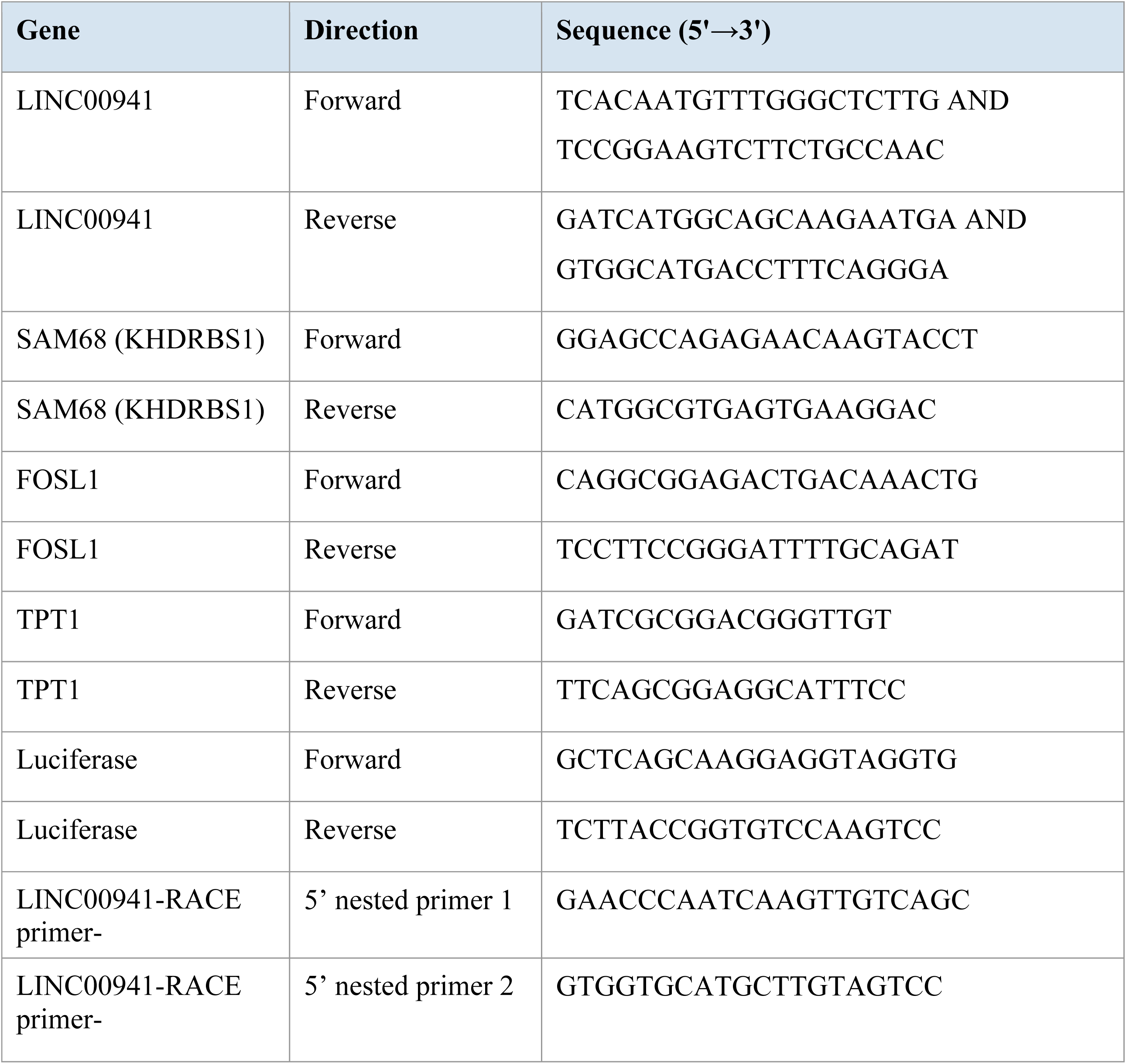
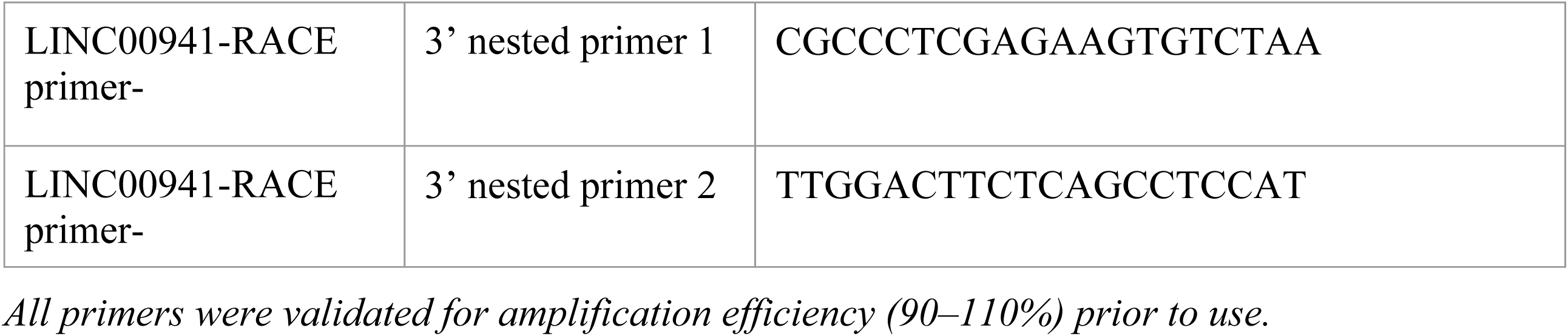
PCR primer sequences used in this study.

**Supplementary Table S3.** Antibodies used in this study.

| Target | Vendor | Dilution (WB / IF / ChIP) |
| --- | --- | --- |
| FOSL1 (FRA-1) | Cell Signaling Technology | 1;1000 (WB) |
| SAM68 (KHDRBS1) | Cell Signaling Technology | 1;1000 (WB),<br>1:100(IF) |
| QKI | Cell Signaling Technology | 1:100 |
| γH2AX (Ser139) | Rockland Immunochemicals | 1:1000 |
| pCHK1 (Ser317) | Cell Signaling Technology | 1:1000 |
| 53BP1 | [PLACEHOLDER] | [PLACEHOLDER] |
| PAR (poly-ADP-ribose) | Cell Signaling Technology | 1:1000 |
| PARP1 | Cell Signaling Technology | 1:1000 |
| β-Actin | BioRad | 1:5000 |
| GAPDH | Cell Signaling Technology | 1:5000 |
| HSP90 | Proteintech | 1:2000 |
| IgG<br>(negative ctrl) | MERCK | ChIP only |

*WB, western blot; IF, immunofluorescence; ChIP, chromatin immunoprecipitation. Dilutions for each application are indicated. — indicates not used for that application.*

**Supplementary Table S4.**
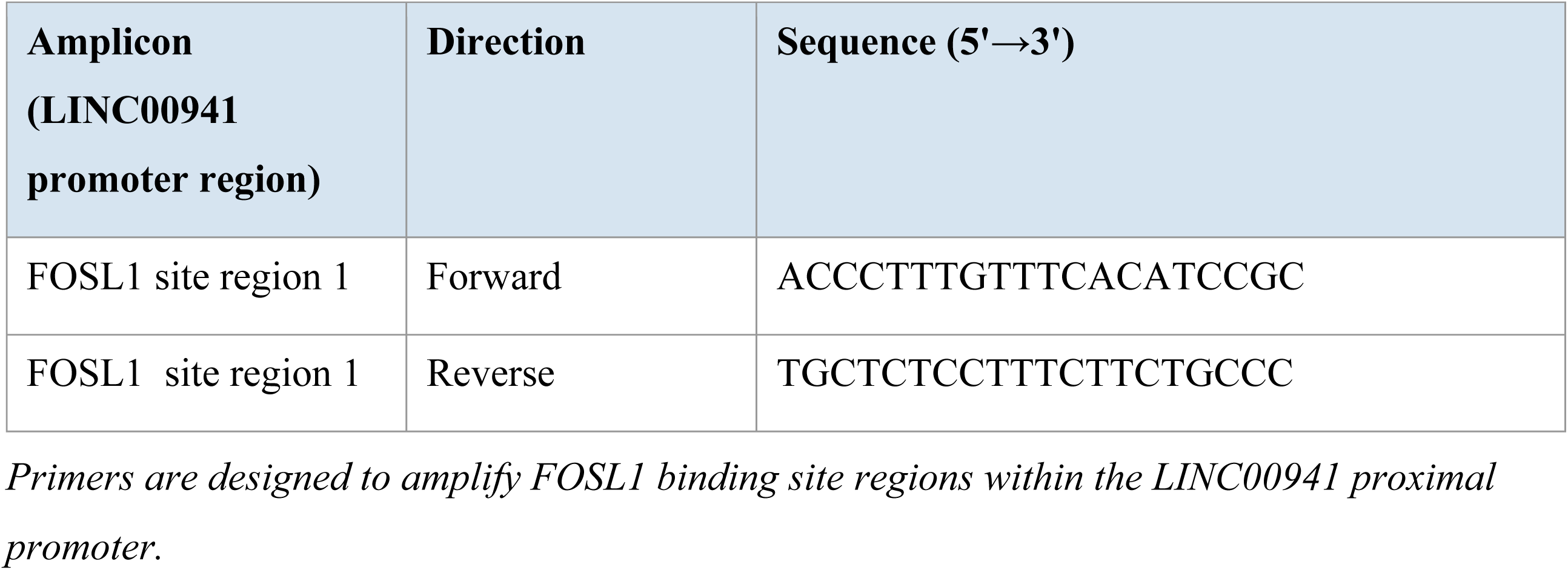
ChIP-qPCR primer sequences for LINC00941 promoter analysis.

**Supplementary Table S5.** KITS and inhibitors.

| Reagent | MAKE | Catalog number |
| --- | --- | --- |
| ChIP Assay Kit | Cell Signaling Technology | 9003 |
| Dual Reporter Assay | Dual-Luciferase® Reporter Assay System | E1910 |
| Cobimetinib(MEKi) | MedChem Express | HY-13064 |
| MG132 | Sigma | M7449 |
| Cyclohexamide | Sigma | C4859 |
| MTT | TCI | 298-93-1 |

**Supplementary Table S6.** Dosing volume of ASOs.

| **Body Weight (g)** | **Volume (uL) BW x 4** |
| --- | --- |
| 15 | 60 |
| 16 | 65 |
| 17 | 70 |
| 18 | 70 |
| 19 | 75 |
| 20 | 80 |
| 21 | 85 |
| 22 | 90 |
| 23 | 90 |
| 24 | 95 |
| 25 | 100 |
| 26 | 105 |
| 27 | 110 |
| 28 | 110 |
| 29 | 115 |

